# Time course of EEG complexity reflects attentional engagement during listening to speech in noise

**DOI:** 10.1101/2023.07.11.548528

**Authors:** Ehsan Eqlimi, Annelies Bockstael, Marc Schönwiesner, Durk Talsma, Dick Botteldooren

## Abstract

Distraction caused by auditory noise poses a considerable challenge to the quality of information encoding in speech comprehension. The aim of this study was to explore the temporal dynamics and complexity of electroencephalography (EEG) microstates in relation to attentional engagement over time, contributing to the understanding of speech perception in noise. We examined three listening conditions: speech perception with background noise, focused attention on the background noise, and intentional disregard of the background noise. Our findings revealed an increase in complexity during the transition of microstates and a slower microstate recurrence when individuals directed their attention to speech compared to tasks without speech. Additionally, a two-stage time course for both microstate complexity and alpha-to-theta power ratio was observed. Specifically, in the early epochs, a lower level was observed, which gradually increased and eventually reached a steady level in the later epochs. The findings suggest that the initial stage is primarily driven by sensory processes and information gathering, while the second stage involves higher-level cognitive engagement, including mnemonic binding and memory encoding.

## Introduction

In the present-day context, the prevalence of distractions poses ongoing challenges for human beings in their daily routines. Among these disruptive elements, diverse forms of noise stand out as a minimally avoidable distraction. This, in turn, emphasizes the substantial influence of these distractions on the auditory context of information encoding, particularly in the domain of attention and speech comprehension (Klatte et al., 2010; Wang et al., 2021a). However, noise exposure does not solely distract; in certain instances, it can positively influence the process of information encoding (Dalton and Behm, 2007; Kiss and Linnell, 2023). A quantifiable and direct estimate of the quality of the information encoding and the impact of noise on cognitive performance is accomplished by analyzing the neural activity signals, surpassing the limitations of traditional methods (e.g. subjective self-report measures and behavioral performance tasks).

The knowledge gained from quantifying cognitive processes and brain states associated with attention and speech comprehension can inform various applications. In educational settings, this knowledge can guide the development of interventions to design more conducive learning environments. For example, educational materials can be tailored to individual cognitive processing abilities, and communication interfaces in artificial intelligence platforms can be improved to optimize students’ attentional engagement. Distraction from content is a key component not only in educational settings during lectures but also in critical contexts such as aviation and medical procedures, requiring understanding and mitigation. Additionally, this understanding can be informed in the design of cognitively-controlled hearing solutions to enhance information processing in noisy environments for hearing-impaired and normal-hearing individuals.

While previous studies (O’sullivan et al., 2015; Han et al., 2019; Geirnaert et al., 2021) have predominantly focused on stimulus reconstruction to address the attentional selection in a cocktail party environment with two speakers, aiming to identify which speaker the listeners are attending to, our study presents a distinct perspective. In lieu of having two distinct speakers, we created a mixture of speech and noise. Participants were instructed to specifically attend to the speech component in the ecologically valid setting (representing conditions that closely resemble everyday listening situations).

Efficiently encoding information during focused speech perception in a noisy setting requires the integrated use of diverse brain resources to facilitate the acquisition and retention of verbal content, which we refer to as "learning from speech". The complexity and existing gaps in our understanding of the underlying mechanisms involved in learning from speech necessitate further investigation. Specifically, delving into the role of attention and memory encoding in this process becomes crucial. By analyzing the dynamic patterns of brain states, particularly through the use of magneto/electroencephalography (M/EEG), it becomes feasible to identify the listener’s attention state. This provides a deeper understanding of the intricate interplay between attention, speech perception, and the learning process.

The brain’s ability to process task-relevant information (target speech) while inhibiting task-irrelevant information (background sound) is known as *attention steering* (Kaya and Elhilali, 2017). Additionally, according to the framework of *predictive coding* (Friston, 2009), the brain encodes the content of speech as prior information and utilizes it to understand and interpret incoming sensory input. Previous studies have extensively discussed the significant role of predictions in shaping perception (De Lange et al., 2018). It has been suggested that conscious perception is linked to the selection of the most probable predictive model (Hohwy et al., 2008; van Kemenade et al., 2020). A simplified Bayesian representation offers an explanation for predictive coding, where the most probable model corresponds to our prior knowledge before listening to the sound, and top-down predictions act as a likelihood (Hohwy et al., 2008). Processing natural continuous speech involves the prediction of forthcoming content. Previous studies have often explored this process by examining the correlation between M/EEG signal (particularly oscillations below 10 Hz) and the acoustic speech envelope (Park et al., 2015; Márton et al., 2019; Koskinen et al., 2020). Indeed, neural entrainment to speech has been proposed as an indicator of predictive coding (Koskinen et al., 2020). Evidence suggests that gamma power, characterized by oscillations above 30 Hz, is influenced by semantic and prediction violation (Penolazzi et al., 2009; Sedley et al., 2016). Additionally, the coupling of theta and gamma oscillations shows potential associations with predictive coding (Hovsepyan et al., 2020). Our earlier research showed compelling evidence that one of the key principal components, obtained by integrating various spectral features including gamma-band activity, exhibits a strong association with predictive coding (Eqlimi et al., 2020, 2021).

*Tulving’s theory* (Tulving, 1972) proposes the existence of two information processing systems that play crucial roles in human memory: *episodic memory*, which pertains to the recollection of temporally dated events, and *semantic memory*. While semantic memory can be activated through language processing, numerous pieces of evidence indicate that non-verbal information processing also contributes to the activation of semantic memory (Bozeat et al., 2000; Ivanova et al., 2021). Furthermore, there is a growing body of research suggesting a temporal and spatial interrelation between these two types of memory (Tulving, 1972; Greenberg and Verfaellie, 2010). The aspect of organizing and ordering information in a temporal sequence has been conceptualized as sequential memory (Tulving et al., 1994). It is important to note that the key distinction between sequential and semantic memories lies in the utilization of prior experiences. While semantic memory retains the actual meaning of information, sequential and episodic memory rely on prior experiences to remember the order in which words or events were presented (Romine and Reynolds, 2004). Numerous neurological studies have been conducted in this area, employing various methodologies. Many of these studies have focused on repeated stimuli and utilized event-related potential (ERP) analysis, such as exploring the correlation between the N400 component and the expectancy of a word (Kutas and Hillyard, 1984). Additionally, investigations have examined the coupling between speech and brain activity during the perception of continuous speech. For instance, it has been proposed that the strong coupling between MEG signals and the speech envelope reflects the predictability of word sequences (Koskinen et al., 2020).

The human brain possesses the remarkable ability to *integrate* information from diverse stimuli presented in its environment. This integration capacity is further complemented by the brain’s capacity to segregate the information into interconnected neural clusters, enabling specialized processing within local functional networks (Mesulam, 1998). In order to support specialized processes, these functional networks undergo rapid reorganization and interconnection, typically within a brief timescale (Bressler, 1995). To understand the dynamics of this neural activity reorganization, it is essential to track the spatio-temporal transient networks that capture the structural and functional architectures.

In the present study, we employed an experimental protocol that builds upon our previous research (Eqlimi et al., 2019, 2020), where we proposed a methodology to assess the functional roles of various frequency-domain characteristics in speech perception under noisy conditions. By implementing this methodology, we showed a significant relationship between EEG and perceptual outcomes (retention of verbal content) (Eqlimi et al., 2020). However, the current study diverges from our previous work by focusing specifically on the investigation of the temporal aspect of brain states during speech-in-noise perception. This distinct focus enables us to address our main research question: how can we observe and study the evolving nature of attentional engagement and memory encoding in the challenging context of speech perception in noisy environments? To achieve this goal, we employed a spatio-temporal analysis by assessing EEG microstates (Lehmann et al., 1987) and proposed a novel methodology to quantify the time-varying complexity of brain states. EEG microstates are quasi-stable scalp topographies that persist within a brief timescale (approximately less than 100 milliseconds) before transitioning swiftly to another topography within the raw and pre-processed multichannel EEG data. The proposed approach involved integrating recurrence quantification analysis (RQA) (Webber Jr and Zbilut, 2005) to investigate the temporal complexity and recurrent patterns of microstates.

The scalp topographies of microstates have been found to exhibit correlations with specific functional magnetic resonance imaging (fMRI) resting state networks (Britz et al., 2010). Spatial cluster analysis enables the extraction of the spatial configuration of microstates. Numerous EEG studies consistently report the presence of four canonical microstate topographical maps, labeled as A, B, C, and D (Koenig et al., 1999; Britz et al., 2010; Milz et al., 2017; Sikka et al., 2020). Changes in microstate statistics, such as duration, occurrence, and coverage, have demonstrated associations with disruptions in mental processes observed in psychiatric disorders like disorders of consciousness (Gui et al., 2020) and Alzheimer’s disease (Tait et al., 2020).

To elucidate the neural mechanisms underlying learning from speech, specifically information encoding and speech comprehension in noisy environments, we employed a combination of listening conditions and tasks. We continuously recorded 64-channel EEG data from N=23 healthy subjects. In the initial stage, we presented several 5-minute sound fragments of lectures covering various topics. These fragments were played in both silence and accompanied by three distinct environmental background sounds, referred to as lecture attended (LA). The purpose of this listening task was to actively engage the subjects and encourage their close attention to the lectures, as their performance on a subsequent exam would be evaluated. Following the lecture stage, we presented multiple excerpts of the background sounds twice. In the first instance, referred to as background attended (BA), participants were instructed to focus their attention on the sounds and attempt to recall the number of salient events, such as a ringing phone. In the second instance, referred to as background unattended (BUA), subjects were instructed to disregard any sounds and not allocate attention to them.

In this paper, our hypothesis revolved around inter-condition and intra-individual differences in terms of EEG microstate properties. Additionally, we anticipated that not only the univariate characteristics, such as microstate duration, might exhibit changes but, more importantly, that the transition between topographies and multivariate characteristics could be influenced by the listening condition. To investigate this, we initially extracted seven microstate topographic classes denoted as A, B, C, C’, D, E, F. Subsequently, we compared the classic microstate features, with a particular focus on microstate duration, and assessed the complexity of the trajectories in the space spanned by these seven microstates using RQA. This comparison was performed among the three different listening tasks: LA, BA, and BUA, with consistent background noise presented in each task. Furthermore, generalized additive mixed modeling (GAMM) was utilized for its capacity to capture individual variations, flexibly model and smooth non-linear trend, enabling us to examine the time-varying changes in microstate complexity and alpha-to-theta power ratio.

## Materials and methods

### Participants

Twenty-three young healthy adults (mean age = 27 ± 3.18 SD, 13 females, 20 right-handed), all English speakers, participated in the experiment. A full battery of audiological tests was conducted, including tonal audiometry, tympanometry, stapedial reflex measurement, speech in noise, and otoacoustic emissions (OAE) with contralateral suppression. No participants were excluded on the basis of this extensive testing of the auditory periphery. All participants provided written informed consent. The study was reviewed and approved by the International Laboratory for Brain, Music and Sound Research (BRAMS) in Montreal, Canada.

### Experimental design

All participants sequentially completed three listening tasks while their 64-channel EEG signals were recorded: 1) attentive listening to English lectures presented in the environmental sound, and 2) attentive, and 3) inattentive listening to fragments of environmental sounds presented alone.

In the first task (referred to as lecture/speech attended, LA), 13 unique lectures were presented, each lasting approximately 5 minutes, with different background sounds at equivalent levels of approximately 63 dBA. The participants were instructed to listen carefully to the lectures and were informed that there would be a written exam after an adequate time interval. In this task, participants actively engaged in learning and retaining the information presented in the lectures, assessing the quality of acquisition and retention of verbal content (referred to as learning from speech). This experimental design ensured a sufficiently long time span between the learning phase and the exam, aiming to minimize the prominence of the last lecture in short-term memory and reduce sequential recall. The lectures covered Belgian cultural and historical topics to ensure minimal prior knowledge in the Canadian test group and to encourage focused attention during the oral presentations. As for the environmental sounds, continuous highway (HW), fluctuating traffic (FT), and multi-talker babble (MT) sounds were used, which introduced informational masking effects, increasing the complexity of the listening conditions and creating a more realistic environment. This type of masking negatively might impact cognitive processing and comprehension of the speech content, even when the speech is audibly clear. During the LA task, lectures were also presented in a silent environment with the addition of pink noise (PK) in the background at an equivalent level of 35 dBA. This inclusion of pink noise created a controlled auditory backdrop, allowing participants to concentrate on the speech content without distractions.

During the second task (referred to as background sound attended, BA) and the third task (referred to as background sound unattended, BUA), fragments of environmental sounds (the same three types used in the first task) were presented without any lectures. The distinction between BA and BUA lies in the participants’ instructions: in BA, they were instructed to pay attention to the sounds, while in BUA, they were instructed to ignore the sounds. These two listening tasks aimed to encompass a wider range of monitored listening conditions and implicitly account for inter-person differences. To promote participants’ attention to environmental sounds, they were asked to memorize the occurrence of salient events, such as phone-ringing, emergency vehicle, and honking car events. Detailed descriptions of stimuli and listening conditions have been provided in (Eqlimi et al., 2020; Eqlimi, 2022). In particular, (Eqlimi et al., 2021) presents an overview of the acoustic characteristics of the presented sounds. The experimental design is displayed in Figure 1.

**FIGURE 1.**
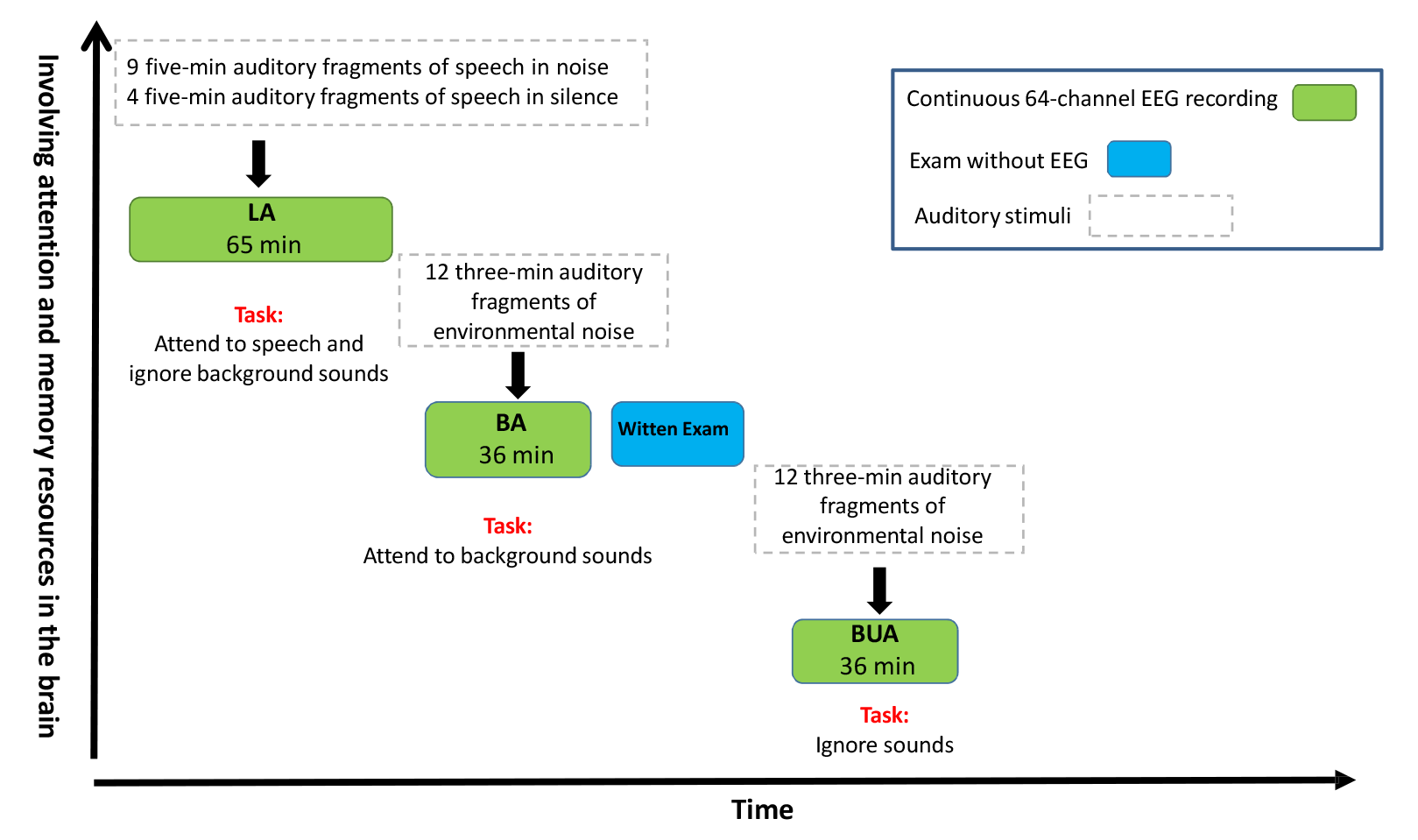
The schematic representation depicts the learning from speech in the noise paradigm employed in our study. Participants underwent sequential exposure to three listening tasks, as indicated by the green blocks from left to right, while their 64-channel EEG signals were recorded. The fugure provides information on the duration, task type, and stimulus type for each condition. Additionally, the blue block represents the exam session (without EEG recording), which focused on the content presented during the lecture/speech attended (LA) condition. The horizontal axis represents the passage of time (from left to right), while the vertical axis (bottom to top) signifies the increasing engagement of task-related cortical activity. For further details, refer to Figure 1 in (Eqlimi et al., 2019).

In this study, there are 10 listening conditions, which are derived from the combinations of three tasks (LA, BUA, BA) and four background sounds (MT, FT, HW, PK): LA-MT, LA-HW, LA-FT, LA-PK, BUA-MT, BUA-FT, BUA-HW, BA-MT, BA-FT, BA-HW. Throughout the paper, the terms "LA-PK" and "LA in silence" are used interchangeably.

### EEG recording and preprocessing

Eyes-open EEG data were collected using a 64-channel EEG recording system (10-20 standard layout, BioSemi System, Amsterdam, NL) with a sampling frequency of 2,048 Hz. Additionally, four extra channels were placed at the outer canthi and below the eyes to record the electrooculogram. The participants were exposed to the three listening tasks mentioned in the previous subsection (LA, BA, and BUA). Each individual had over 2 hours of EEG data recorded (approximately 65 minutes for LA, 36 minutes for BA, and 36 minutes for BUA).

EEG preprocessing was conducted based on the EEGlab version 13.1.1b (Delorme and Makeig, 2004) and FASTER (Nolan et al., 2010) toolboxes. Preprocessing was performed for each individual listening task, involving the continuous EEG signal that encompasses all fragments presented during each task. The EEG data were first offline referenced to the nose electrode, which functioned as the external channel, and resampled at a frequency of 512 Hz. Subsequently, the data were band-pass filtered between 0.5 and 135 Hz using a 3, 380*^th^* order FIR band-pass filter with a Hamming windowed sinc. Eye-related artifacts were visually identified by inspecting the independent components, and subse-quently removed from the EEG data (for further details, see (Eqlimi et al., 2020)).

The preprocessed continuous EEG data for each individual listening task was split into fragments, corresponding to the 3-minute (BA and BUA) or 5-minute (LA) exposures. The sound signals were recorded concurrently with the EEG data using dedicated external channels integrated into the EEG setup. To compensate for any potential delays, the synchronization technique described in our previous study (Eqlimi et al., 2020) was employed to ensure accurate synchronization between the auditory stimuli and the EEG measurements. The synchronized EEG signal per fragment was subjected to channel cleaning utilizing the FASTER algorithm (Nolan et al., 2010), with a Z threshold of 3, which represents a measure of standard deviation from the mean, to eliminate bad channels. The eliminated channel signals were subsequently interpolated using a weighted neighbor approach.

To ensure consistency in main feature calculation (TFR, microstate, and complexity), the initial 3-minute segment of data from the BA, BUA, and LA tasks was utilized, minimizing bias and facilitating fair feature comparisons. Specific preprocessing steps, including bandwidth filtering and common average re-referencing, were performed for the main analysis of EEG data, with detailed explanations provided in their respective sections. Furthermore, given the extended duration of our experiment, we conducted assessments to quantify potential fatigue-related effects and movement artifacts. The supplementary findings indicated no considerable differences in the occurrence of eye, mus-cle, heart, and channel artifacts among the three consecutive listening tasks. This observation suggests effective mitigation of unwanted fatigue-related effects, which could potentially reduce the quality of the data over the course of the experiment.

### Time-frequency analysis

First, the preprocessed EEG signals within each fragment were segmented into 1-second epochs. Time-frequency (TF) analysis was subsequently performed on each epoch using the multi-taper method fast Fourier transform (“mtmfft”) with the Fieldtrip toolbox (Oostenveld et al., 2011), resulting in complex-valued TF planes for each epoch denoted as **F**. The power of each TF point, denoted as **P**_EEG_, was then computed for all individual epochs using the following equation:

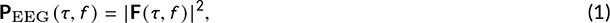

where *τ* and *f* denote the time and frequency bins, respectively.

The EEG TF representation was estimated over a frequency range of 1 − 45 Hz, with 1 Hz steps, for every 100 milliseconds. This estimation was performed using a Hanning taper with a 50% overlap and a spectral smoothing of 1 Hz. For each epoch, the average EEG power (averaged over time) was calculated for the *θ* (4 − 8 Hz) and *α* (8 − 12 Hz) frequency bands. As the data were paired (each subject measured multiple times), no normalization, such as normalizing to total power or baseline, was applied. The epoch-wise *α* /*θ* power ratio, averaged over all channels, was then used for statistical modeling (refer to subsection).

### Microstate analysis

The EEG topographical maps exhibit a quasi-stable state for a sub-second time period before rapidly transitioning into different topographical patterns (Lehmann, 1971). These quasi-stable topographical maps, referred to as EEG microstates, can be extracted by performing a clustering analysis on all conditions, as conducted in this study, or on each condition individually. Extracting microstates from pooled data across all conditions provides the advantage of incorporating contributions from all explicit listening conditions into the clustering analysis, similar to the method used for computing principal components in (Eqlimi et al., 2020). This pooling approach facilitates the identification of microstates that are consistently expressed under specific listening conditions.

The template microstate maps were identified as follows. Firstly, the EEG data were band-pass filtered between 1 − 45 Hz and re-referenced to the common average reference. The aggregated within-subject data was created, containing observations of all listening tasks (BA**+**BUA**+**LA, • **+** • stands for concatenation of two dataset) from the subject. The EEG signals were aggregated by concatenating the individual EEG matrices along the time axis. The resulting aggregated matrix had dimensions of 64× (12*N*_3min_), 64× (12*N*_3min_), and 64× (13*N*_5min_), representing 12 BUA fragments, 12 BA fragments, and 13 LA fragments of preprocessed EEG matrices, respectively. Here, *N*_3min_ represents the number of time samples in the BA and BUA EEG signals, and *N*_5min_ represents the number of time samples in the LA EEG signals. The total dimensions of the aggregated matrix, **S**, were 64 × (12*N*_3min_ + 12*N*_3min_ + 13*N*_5min_).

Single-subject microstate topographies, also known as personalized templates, were extracted using a modified k-means clustering method (Pascual-Marqui et al., 1995) (see supplementary text, section 1) 40 repetitions, disregarding map polarity (topographical maps are polarity invariant, i.e., **a***_q_* = −**a***_q_*; see supplementary text, section 1) and at the time of global field power (GFP) peaks, based on (Poulsen et al., 2018). The extraction was performed on the aggregated matrix, **S**.

The optimal number of brain states was determined to be seven based on the Krzanowski-Lai (KL) criterion(Krzanowski and Lai, 1988). As individual microstate maps have no inherent order, a permutation method was employed to compute the grand average microstate maps across subjects. This permutation involved rearranging the extracted topographic maps within each subject to maximize commonality across cases, following the algorithm proposed in (Koenig et al., 1999).

Two approaches can be adopted to identify the sequence of microstates. In the first approach, assuming microstate discreteness, the label of the most similar microstate map is assigned to the momentary topographical maps, generating a univariate microstate sequence (e.g., 𝕊 = {A, B, C, …, F, B, D}) is generated for each fragment of EEG data (subsection). In the second approach (used in the subsection of), the original dissimilarity values of the microstate maps with the momentary topographical maps are used. Instead of a univariate sequence, a microstate matrix **M** ∈ ℝ^7×*n*^_*t*_ is computed, where 7 is the number of microstate classes, and *n_t_* is the number of time intervals. A similar terminology and approach have been utilized in (Pascual-Marqui et al., 2014).

### Back-fitting microstates to EEG samples

Following the alignment of the individual microstate maps and computing the grand average microstate maps, we employed the global map dissimilarity (GMD) (Murray et al., 2008) measure to determine the signal strength-invariant distance between the template microstate topographies and the instantaneous EEG topography. Alternative strength-invariant criteria, such as cosine distance, can also be utilized. The GMD quantifies the topographic disparities between two potential maps, denoted as **u** and **v**, providing a measure of the dissimilarity in their spatial distribution. The GMD can be calculated using the following formula:

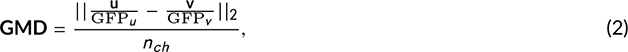

where **u** and **v** represent either two instantaneous topographical EEG maps (in subsection to identify microstate maps) or one instantaneous topographical EEG map and a template microstate (here, for the back-fitting process), *n_ch_*is the total number of EEG channels, and **GFP***_v_* refers to the spatial standard deviation (*ℓ*_1_-norm) or the total energy (*ℓ*_2_-norm) in the topography over time as follows:

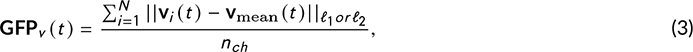

where **v***_i_* (*t*) is the *i ^th^* EEG channel signal at time *t*, **v**_mean_ (*t*) is the averaged EEG signal across channels at the time *t*. Both *ℓ*_1_ and *ℓ*_2_-norm can be used to calculate GFP, as they provide similar microstate results (Milz et al., 2016). In this paper, we used *ℓ*_2_-norm.

To implement the winner-take-all hypothesis, we assigned each original topography at every time instant to a template microstate map based on the minimum GMD values. This allowed us to label the EEG topography at each time point *t* according to the most similar template map. Next, we applied temporal smoothing to the microstate label sequences, disregarding segments with a duration of less than 30 milliseconds (Poulsen et al., 2018). The resulting microstate sequences were then used to calculate important microstate statistics, including the mean duration (the average duration that a given microstate occurs), occurrence (the number of times that a specific microstate occurs in a second), and coverage (the percentage of time spent within a microstate).

We opted to apply back-fitting and calculate microstate statistics on the data epoched into 100 millisecond segments to ensure consistency with the extraction of complexity features, which were also computed at the same 100 millisecond intervals. Furthermore, we included the results from the analysis of the non-epoched data in the results section.

### Microstate complexity analysis

The Markovian transition analysis, also known as syntax analysis, is a commonly used method to examine microstate transitions. It centers the current microstate to estimate the probabilities of transitions. However, it assumes that these transition probabilities are stationary, even though there is evidence suggesting that microstate transitioning exhibits non-stationary behavior (Van de Ville et al., 2010; von Wegner et al., 2017). Furthermore, due to the long-range dependence observed in microstate sequences (Van de Ville et al., 2010), a Markovian model is inadequate in capturing this long-range memory.

In this study, we introduce a new approach to analyze the complexity of microstate transitions that leverages recurrence quantification analysis (RQA) (Webber and Marwan, 2015). Remarkably, this approach is applied unprecedentedly within the microstate context, addressing the limitations of previous approaches and capturing the non-stationary nature of microstate transitions.

Unlike previous EEG-related studies that rely on the embedding technique to construct the phase space (Gruszczyńska et al., 2019; Fan and Chou, 2018), our approach utilizes the microstate GMD time series. These time series consist of distances between the EEG topography per 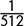 second segment and each of the seven microstate topographies. To create the microstate-based phase space, we averaged the GMD time series over a moving window of length 100 milliseconds. As a result, the microstate-based phase space captures microstate transitions with a final temporal resolution of 100 milliseconds, represented in a seven-dimensional domain. To generate thresholded recurrence plots (RPs), we compute the pairwise Euclidean distance between the GMD vectors (treated as phase vectors) according to the method proposed in (Marwan et al., 2007).

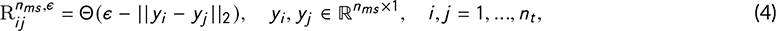

where *n_ms_*represents the number of microstates, which corresponds to the dimension of the phase space, *n_t_*is the number of time points (with the step of 100 milliseconds), Θ(•) denotes the Heaviside function, *ɛ* is a threshold distance, *y_i_*, and *y_j_* are the GMD values at time *i* and *j*, respectively.

Aiming to maintain a consistent number of recurrence connections across all individuals and time windows, a proportional thresholding approach was adopted. The goal was to preserve a fixed percentage, denoted as *e*%, of the smallest distance connections. This approach aimed to ensure a comparable level of recurrence connectivity throughout the dataset.

Two windowing approaches can be employed to estimate time-dependent RQA. In the first approach, the GMD time series is segmented into windows of size *l_w_* (here, *l_w_* = 10 corresponding to 1 second), and the recurrence plot and RQA measures are computed for each window individually. This approach allows for the examination of temporal variations in RQA measures within distinct time intervals. In the second approach, the recurrence plot is portioned into the windows of size *l_w_* that extend along the line of interest (LOI), and within each window, the RQA features are computed. The second approach was chosen for this study because, unlike the first approach, it determines the threshold based on the overall distribution of distances. Specifically, the threshold is set so that a certain percentage (*e*%, with a 30*th* percentile chosen in this paper) of the smallest distance connections are retained. This threshold is then applied to each time window consistently. However, it should be noted that considering different thresholds in the first approach could be valuable for detecting nonstationarities within the data.

**FIGURE 2.**
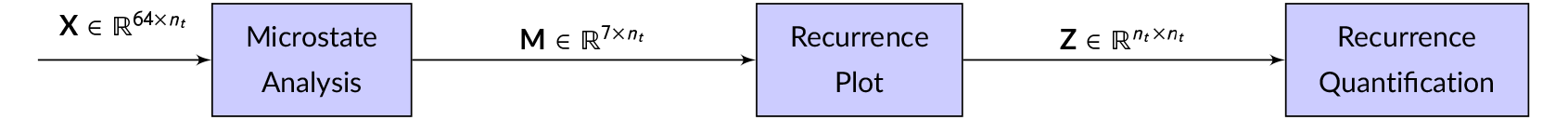
Block diagram illustrating the proposed analysis for assessing the complexity of EEG signals. The diagram shows the flow of information from the 64-channel EEG signals (**X**) to the Global Map Dissimilarity (GMD) matrix (**M**), quantifying the dissimilarity between EEG samples and the pre-extracted microstate topographies, and further to the two-dimensional representation of microstate recurrence (**Z**). The analysis considers 7 microstate classes, with the parameter *n_t_* representing the number of time points included.

Since the RQA features rely on histograms, a window size of *l_w_* = 10 was used to ensure an adequate number of samples for feature quantification. Multiple features were estimated for each 1-second recurrence plot. Among these features, this study places particular emphasis on recurrence time (Ngamga et al., 2012), which measures how frequently a small region in the phase space (in this case, the EEG microstate space) is visited. Two types of recurrence time statistics have been developed based on the scaling law proposed in (Gao, 1999). The first type is the Poincare recurrence time, also known as the recurrence time of the first type 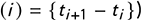, which represents the occurrence of true recurrence points. The second type is the recurrence time of the second type 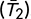, which represents the occurrence of sojourn recurrence points. For discrete maps, 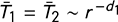, where *r* is the radius of the neighborhood and *d*_1_ is the information dimension. In this study, we will refer to the recurrence time as T_2_. Practically, T_2_ can be estimated based on the histogram of white (representing **R***_ij_* = 0) vertical lines in the recurrence plot, as follows:

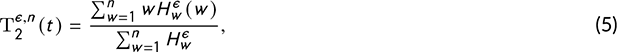

where *n* represents the number of time points in each sub-matrix of the recurrence plot, *w* and *H_w_* are the length and histogram of white vertical lines, respectively. T_2_ corresponds to the period of periodic signals and is associated with the Kolmogorov entropy of chaotic signals (Gao and Cai, 2000). This measure has been proposed for detecting epileptic seizures using EEG (Gao and Hu, 2013). Figure 3 demonstrates the method to calculate T_2_.

**FIGURE 3.**
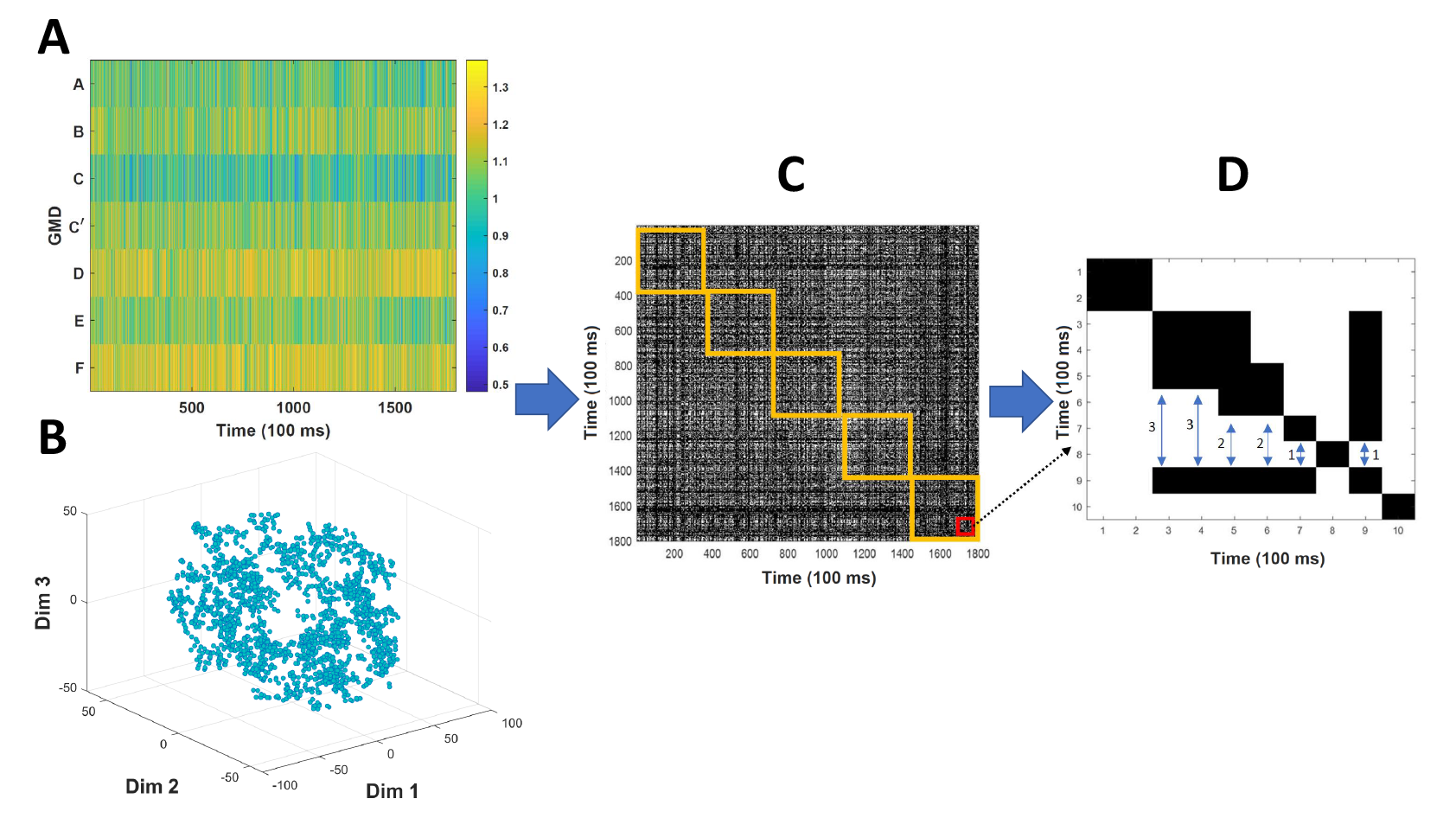
An illustrative block-diagram to demonstrate how to calculate the recurrence time (T_2_). **A**) The global map dissimilarity matrix represents a 7-dimensional microstate space. **B**) The microstate space is visualized in a 3-dimensional representation using t-distributed stochastic neighbor embedding (t-SNE), which is not part of the main analysis. **C**) A binary recurrence plot (RP) is extracted from the 7-dimensional microstate space, where black dots indicate **R***ij* = 1 or recurrence points, and white dots indicate **R***_ij_* = 0. The yellow-colored squares indicate the sub-RPs used for estimating complexity features. **D**) To calculate T_2_, the number of vertical white lines is counted before a recurrence point or black dot. In this example, there are 2 vertical white lines of length 1, 2 vertical white lines of length 2, and 2 vertical white lines of length 3. Applying equation 5, we find that 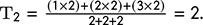

### Statistical analysis

Linear mixed-effect modeling was applied using the LME4 package of the statistical software R (Bates et al., 2015). Two distinct statistical models were performed: 1) for each background sound, including observations from all listen-ing tasks for the *k ^th^* background sound, and (2) for each listening task, including observations from all background sounds for the *i ^th^* listening task. In the first model, referred to as the within-background model, the time-averaged microstate features were used as separate response variables in linear mixed-effects models. The fixed categorical factor included the *listening task* (BA, BUA, and LA), while the random categorical factor accounted for *participant* vari-ability (considering between-subject differences). The second model, referred to as the within-task model, is similar to the first model, with the difference that the fixed categorical factor is based on *background sound* (MT, FT, and HW) rather than the *listening task*. The formulas for the two models are as follows:

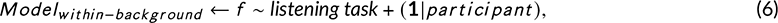

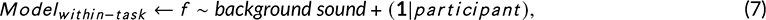

where *f* is the time-averaged microstate feature (e.g. mean duration of microstate A), the symbol ∼ implies that the left term is modeled as a function of right terms, and random terms are expressed in (**1**|*participant*).

The *χ*^2^ difference test can be utilized to assess the significance of differences in the Akaike information criterion (AIC) when adding or eliminating model parameters. In our study, we employed this test to compare the models that included the fixed variable of interest, either the *listening task* or the *background sound* factor, with the models that did not include these factors. Pairwise Tukey *post hoc* comparisons with Bonferroni–Holm correction were performed to determine the significance of differences between the listening tasks for each background sound (based on the results of the within-background model) and between background sounds for each listening task (based on the results of the within-task model).

### Modeling time-varying neural features

To account for the temporal autocorrelational structure and inter-subject variability in the neural time series obtained in previous subsections, we employed a generalized additive mixed model (GAMM) (Wood, 2006). GAMM extends the capabilities of the generalized linear mixed model (GLMM) by incorporating unknown and non-linear smooth functions of the predictors. By incorporating both the random factor of *participant* and the non-linear components, GAMM enables us to capture complex temporal dependencies and non-linear relationships in the data.

The GAMM can be represented in the following general form:

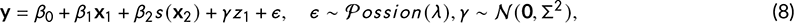

where **y** is the response variable, the contributions of *x*_1_ and *x*_2_ are linear and non-linear, respectively. This means that the contribution to the linear predictor includes *x*_1_ and a function *s* of *x*_2_ (smooth term). Adding the random effect, *z*_1_, leads to fitting a mixed-effect model.

In our study, three variables were used to model the response variable: 1) a categorical variable, *listening condition*, which encompasses 10 possible values combining the listening tasks. i.e., LA, BA, and BUA and background sounds. i.e., MT, HW, FT, and PK (only for LA), 2) a continuous variable, *time*, and 3) a categorical variable, *participant*, which is treated as a random effect. Therefore, the general form of our model is the following:

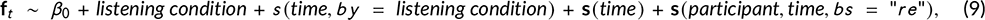

where **f***_t_* is the time-varying neural feature (e.g, the microstate recurrence time, T_2_), *s* (•), *by* = •, and *bs* = "*re*" represent the smooth function, interaction, and the random effect, respectively. Random smooth functions adjust the slope of the trend of the numeric predictor (*time*) using **s**(*participant*, *time*, *bs* = "*r e*") term. It is important to note that entirely separate smooth functions are fitted for each listening condition. The GAMM analysis was performed using the MVGC package in the statistical software R (Wood, 2006).

## Results

### Microstate topographies

The median optimal number of microstate templates, determined using the Krzanowski-Lai criterion, was found to be *k* = 7 across all subjects, within a range of 4 − 15 clusters. Figure 4 displays the topographical maps of each of these seven microstates, which exhibit a strong resemblance to the classical microstate maps (A-F and C^′^) (Custo et al., 2017; Michel and Koenig, 2018; Tarailis et al., 2023). These classical microstate maps have been shown to align with observations in fMRI and simultaneous EEG-fMRI studies (Britz et al., 2010).

**FIGURE 4.**
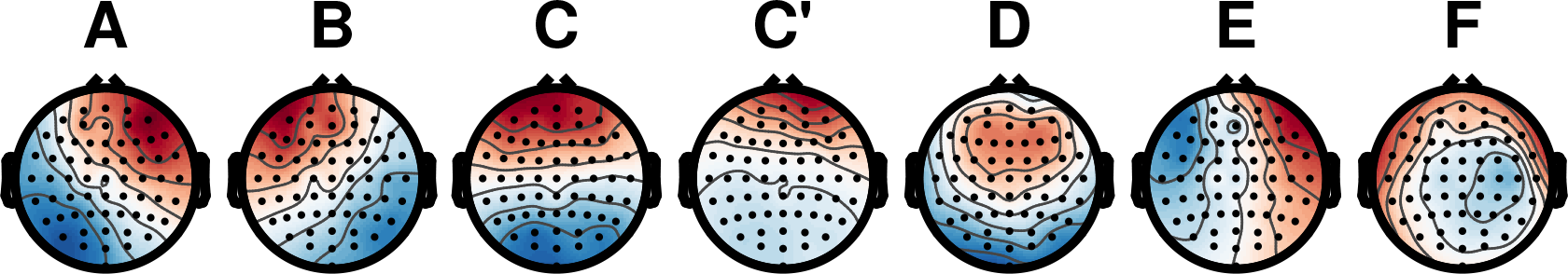
Topographical maps of the seven prominent microstates were extracted and sorted across 23 subjects. The EEG signals were initially aggregated by concatenating the individual EEG matrices along the time axis. Subsequently, microstates were identified within each subject, and the average of these microstates was calculated across all subjects.

Specifically, microstates A-D are associated with resting state networks (RSNs) as follows: auditory (A), visual (B), saliency (C), and attention/working memory (D). However, it is important to note that certain studies, which we will discuss further in the discussion section, argue against the existence of a simple relationship between microstates and RSNs. The explanations for microstates C^′^, E, and F can be found in (Michel and Koenig, 2018). It should be noted that microstate C^′^ in the current study corresponds to microstate C^′^ in the works of (Michel and Koenig, 2018; Jabès et al., 2021), microstate F in (Custo et al., 2017), microstate 3 in (Britz et al., 2010), and microstate E in (Zanesco et al., 2020). Additional information and detailed results regarding microstate topographies can be found in the supplementary text.

### Microstate statistics

The temporal dynamics of microstates can be quantified in terms of the basic features, namely, duration, occurrence, and coverage for each microstate. Here, our emphasis was on the microstate duration. First, the average microstate duration over time (time-averaged feature, *f*) in each microstate was compared between different conditions using the linear mixed-effect modeling as described in subsection.

We revealed the *listening task* factor (LA, BA, and BUA) improves the predictability of the mean duration of microstates (except microstate C^′^ for all the sounds and microstate E for the highway sound) compared to the model without fixed predictor for all the background sounds (Table 1). On the contrary, there was no significant improvement found by adding the *background sound term* (MT, FT, HW). Tukey *post hoc* comparison showed a significant decrease in mean microstate duration in BUA and LA compared to BA, averaged over all microstates (BA > BUA, *p* < 10^−7^ and BA > LA, *p* < 10^−7^). Note that this comparison was performed taking into account all the environmental sounds and ignoring the fragments presented in silence in the LA. Similar significant differences were found for the mean microstate occurrence within the background sounds and between the listening conditions, i.e. BA exhibits a minimal microstate occurrence.

**TABLE 1.**
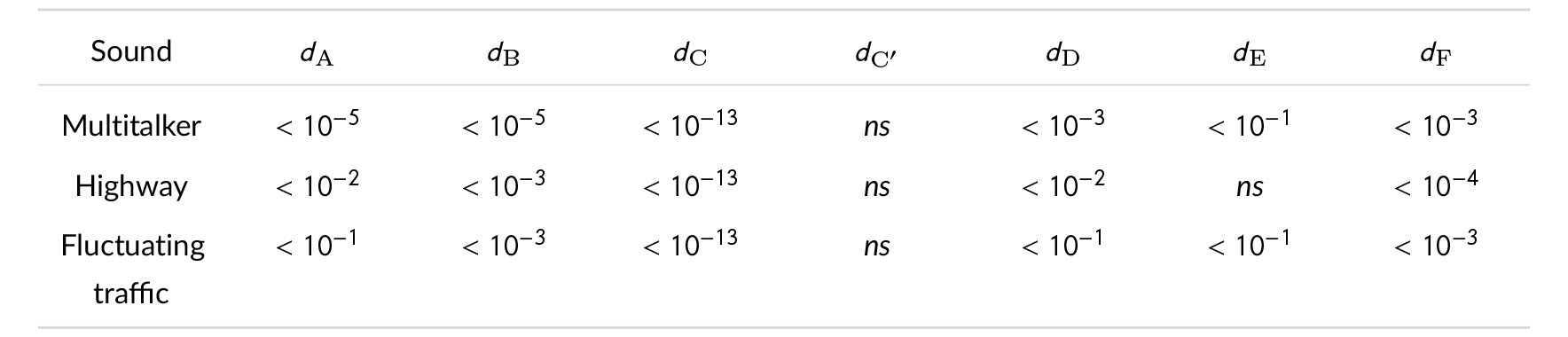
Effect of listening task on the predictability of microstate duration; The table presents the p-values resulting from the *χ*^2^ difference test between the baseline model and the model presented in equation 6 for each background sound. A non-significant effect (*p* > 0.05) is denoted as "ns". *d*_A-F,C_′: mean duration of microstates A-F&C^′^.

| Sound | $d_A$ | $d_B$ | $d_C$ | $d_{C'}$ | $d_D$ | $d_E$ | $d_F$ |
| --- | --- | --- | --- | --- | --- | --- | --- |
| Multitalker | $< 10^{-5}$ | $< 10^{-5}$ | $< 10^{-13}$ | ns | $< 10^{-3}$ | $< 10^{-1}$ | $< 10^{-3}$ |
| Highway | $< 10^{-2}$ | $< 10^{-3}$ | $< 10^{-13}$ | ns | $< 10^{-2}$ | ns | $< 10^{-4}$ |
| Fluctuating traffic | $< 10^{-1}$ | $< 10^{-3}$ | $< 10^{-13}$ | ns | $< 10^{-1}$ | $< 10^{-1}$ | $< 10^{-3}$ |

Figure 5 represents the violin plots of mean duration for each microstate along the conditions in which the multi-talker sound was presented as background. Stars denote the pairwise significant differences between the conditions based on the Tukey *post hoc* analysis. These results are suggestive of (I) shorter microstate A in the LA, (II) longer microstate B in the LA, (III) shorter microstate C in the BUA, (IV) shorter microstate D in the LA (only compared to the BUA), (V) longer microstate E in the LA (only compared to the BUA), and (VI) longer microstate F in the LA. The mean duration of microstate C^′^ was not significantly different between the tasks.

**FIGURE 5.**
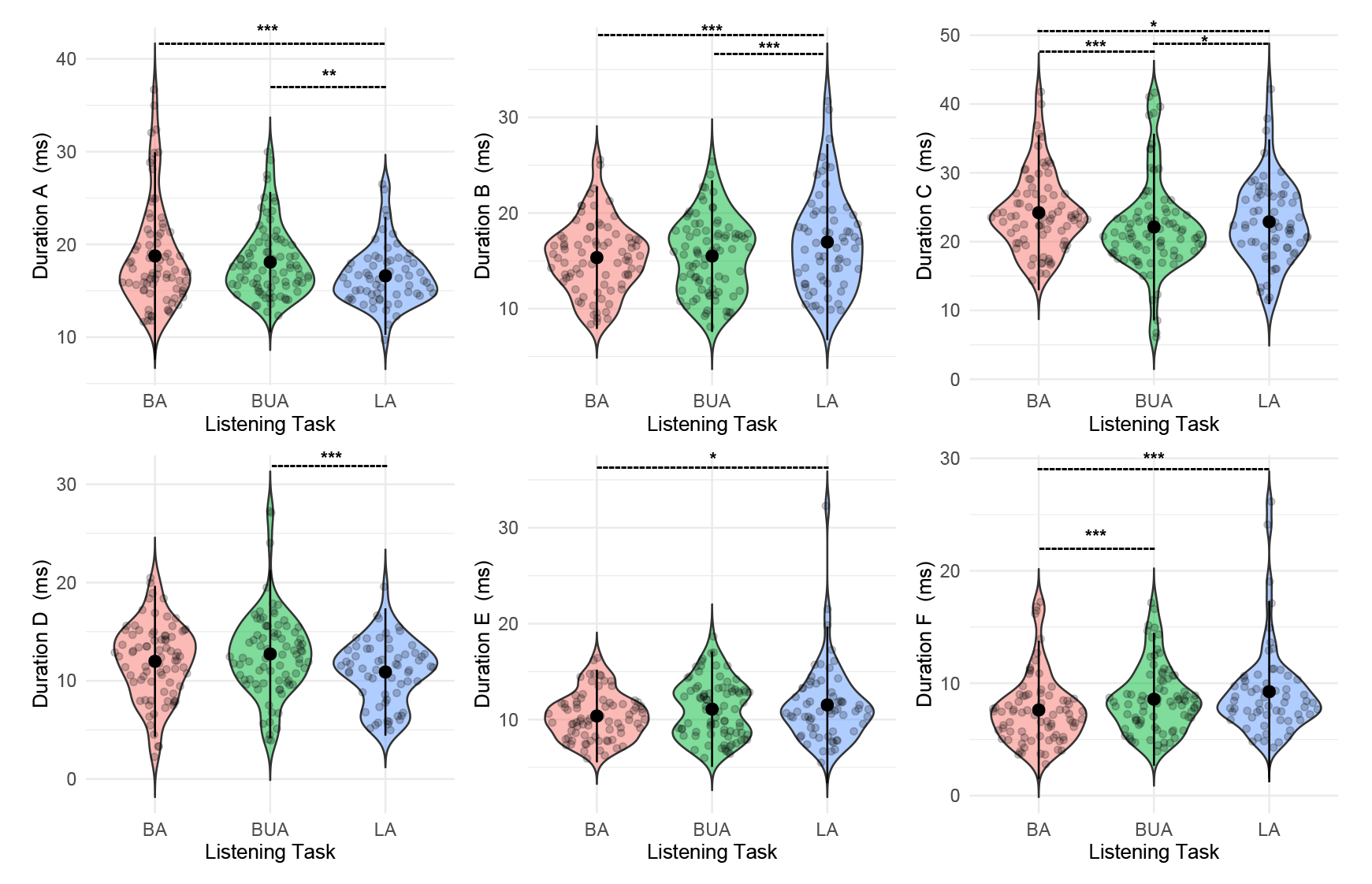
Mean duration of microstates with significant differences for the three listening tasks (BA, BUA, and LA) in the multi-talker (MT) sound. The data points show values for each participant per fragment. Stars denote *p*-values obtained based on the linear mixed-effect model with *participant* as a random factor using *post hoc* pairwise comparisons with Bonferroni correction: ∗*p* < 0.05, ∗ ∗ *p* < .01, ∗ ∗ ∗*p* < 0.001.

The similar trends are observed in the *post hoc* pairwise comparisons for the highway and fluctuating sounds. For the highway sound, the results are as follows: [*d*_A_; BA>LA, *p* < 0.01], [*d*_B_; LA>BA, *p* < 0.001 & LA>BUA, *p* < 0.001], [*d*_C_; BA>LA, *p* < 0.001 & LA>BUA, *p* < 0.05], [*d*_D_; LA>BA, *p* < 0.05 & LA>BUA, *p* < 0.01], and [*d*_F_; LA>BA, *p* < 0.001 & BUA>BA, *p* < 0.001], where *d*_A-F_ denotes mean duration of microstates A-F.

For the fluctuating traffic sound, the results as follows: [*d*_A_; BA>LA, *p* < 0.05], [*d*_B_; LA>BA, *p* < 0.001 & LA>BUA, *p* < 0.01], [*d*_C_; BA>LA, *p* < 0.001 & LA>BUA, *p* < 0.05], [*d*_D_; LA<BUA, *p* < 0.01], and [*d*_F_; LA>BA, *p* < 0.001 & BUA>BA, *p* < 0.01].

### Microstate complexity

The results of quantifying the transition complexity of microstates are presented in two subsections. First, the mean values, i.e., the time-averaged values between different listening tasks, are compared by linear mixed-effect model, and then in the second subsection, the time-varying complexity of microstates are modeled and compared between the conditions based on GAMM. The six complexity features have been assessed in this study: recurrence rate, determinism, entropy, laminarity, trapping time, and recurrence time. The details of the calculation of these features are explained in the supplementary text, section 2.

### Mean complexity

Among the complexity features, for the trapping time (in the MT sound) and the recurrence time (in all background sounds), we found a significant improving effect by including the listening task compared to the model without fixed predictor (Table 2).

**TABLE 2.**
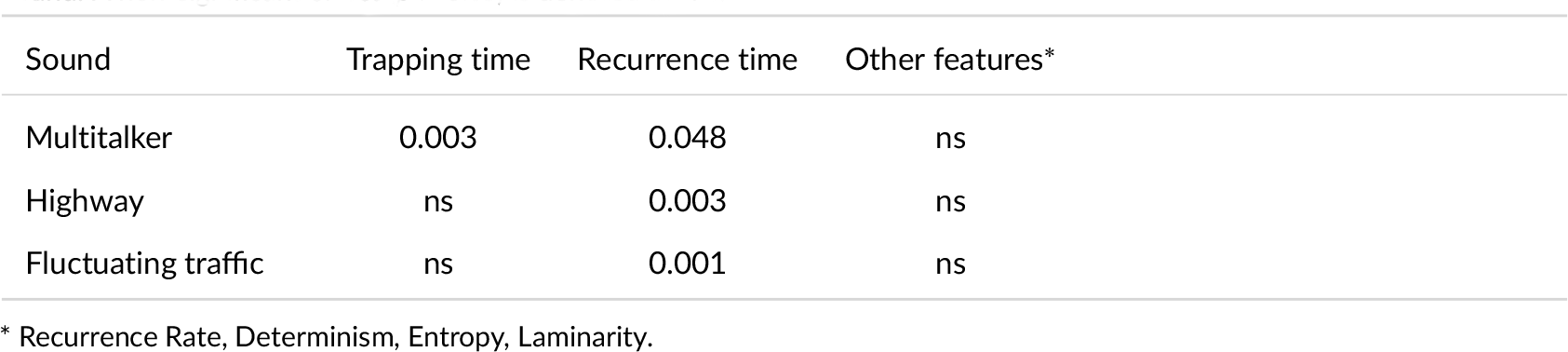
Effect of listening task on the predictability of microstate complexity features; The table presents the p-values resulting from the *χ*^2^ difference test between the baseline model and the model presented in equation 6 for each background sound. A non-significant effect (*p* > 0.05) is denoted as "ns."

| Sound | Trapping time | Recurrence time | Other features* |
| --- | --- | --- | --- |
| Multitalker | 0.003 | 0.048 | ns |
| Highway | ns | 0.003 | ns |
| Fluctuating traffic | ns | 0.001 | ns |

\* Recurrence Rate, Determinism, Entropy, Laminarity.

Figure 6 represents the violin plots of mean complexity measures of microstate along the conditions in which the multi-talker sound was presented as background. Stars denote the pairwise significant differences between the conditions based on the Tukey *post hoc* analysis.

**FIGURE 6.**
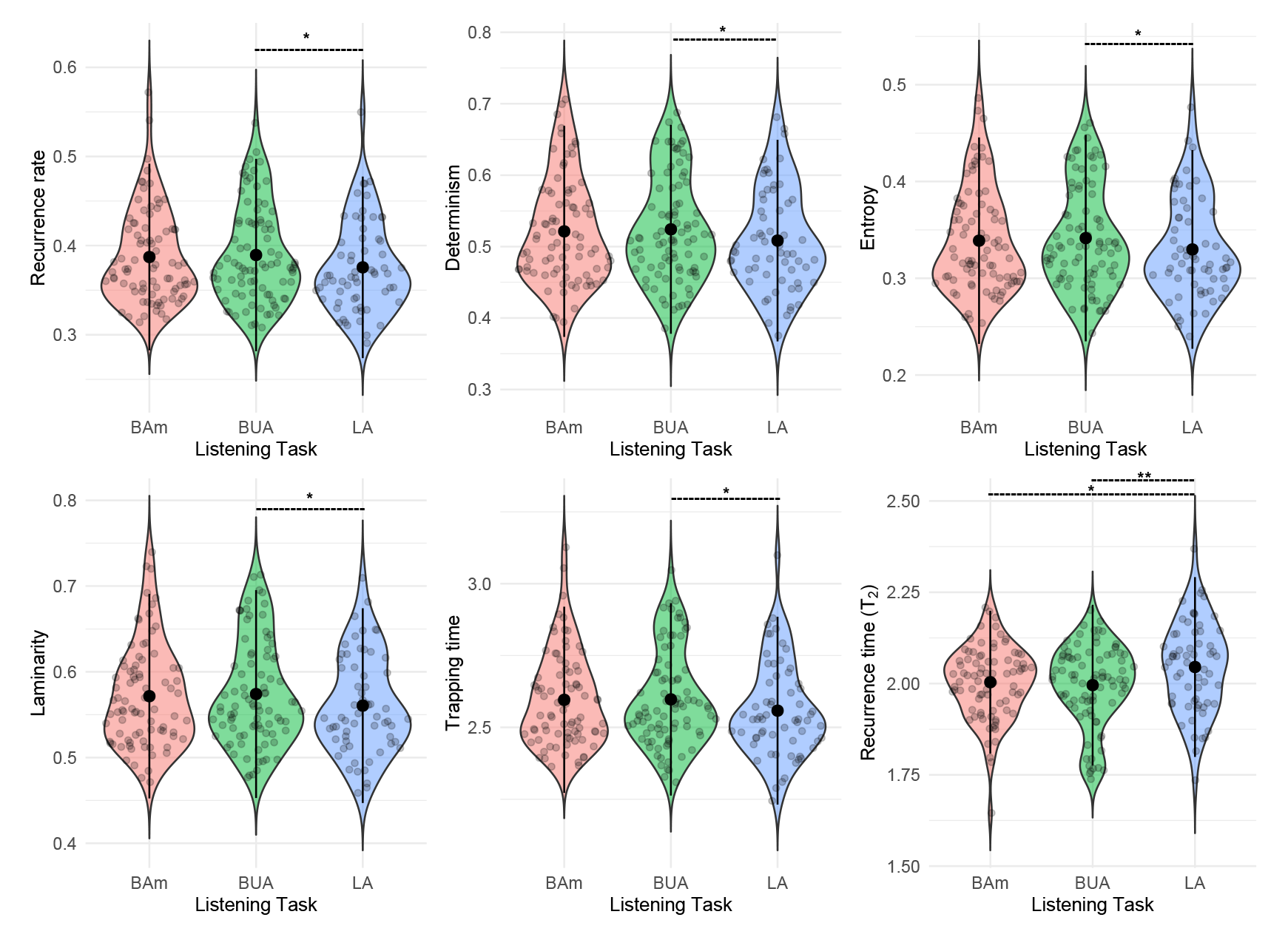
Mean (time-averaged) complexity measures of microstate for trajectories in three listening tasks (BA, BUA, and LA) in the multi-talker (MT) sound. The data points show values for each participant per fragment. Stars denote *p*-values obtained based on the linear mixed-effect model with *participant* as a random factor using *post hoc* pairwise comparisons with Bonferroni correction: ∗*p* < 0.05, ∗ ∗ *p* < .01, ∗ ∗ ∗*p* < 0.001.

The Tukey *post hoc* comparison showed the least complex microstate transitions occur in BUA. The complexity feature, T_2_ (recurrence time), is significantly increased in LA. Furthermore, the features trapping time and laminarity significantly decrease in LA. The laminarity tries to quantify the intermittency of microstate transitioning, namely, the irregular alternation of phases in the recurrence plot. Trapping time is associated with the predictability time, namely, how long the system remains in a specific state. Therefore, the second type of recurrence points, namely, hidden states, remains longer in the LA compared to BA and BUA.

In the *post hoc* pairwise comparisons for the highway and fluctuating traffic sounds, the feature T_2_ was only significantly different between the tasks: [highway, T_2_; LA>BA, *p* < 0.01 & BUA>BA, *p* < 0.05] and [fluctuating traffic, T_2_; BUA<BA, *p* < 0.001].

### Effect of time: Within-task fragment order

As mentioned, in this study, 10 different listening conditions are presented to the participants. Each listening condition is determined by the type of listening task, i.e., LA, BA, and BUA, and the type of background noise, i.e., PK (only in LA), MT, HW, and FT. In this subsection, we aim to study the effect of the order of the presented fragments on T_2_, regardless of which background sound is used in each task. In the BA and BUA tasks, 12 fragments of background sounds were presented, and in the LA task, 13 fragments of speech plus background sounds were presented. We first grouped each listening task into 4 groups based on the order of presenting the fragments. *Fragment Order* was then used as a categorical variable to investigate the effect of time on T_2_ using the mixed-effect modeling as follows:

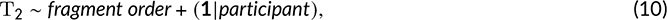

where T_2_ is the time-averaged recurrence time, *participant* is the random factor, *fragment order* is the fixed factor that has 4 possible outcomes: *O*_1_ (fragments 1, 2, and 3), *O*_2_ (fragments 4, 5, and 6), *O*_3_ (fragments 7, 8, and 9), and *O*_4_ (fragments 10, 11, 12 and 13). This regression was done separately for all observations in each listening task. Tukey *post-hoc* multiple comparisons were then performed to test for the significance of T_2_ changes across the fragment order. In the LA and BUA listening tasks, no significant difference (with the level of significance of 0.05) was found between each pair of fragment orders, meaning no decreasing or increasing trend was observed in T_2_ due to the order of presentation of the fragments. In the BA listening task, there was only one significant difference between *O*_3_ and *O*_4_, which showed that T_2_ in *O*_4_ (the last fragment) increases significantly (adjusted *p*-value= 0.0074) compared to *O*_3_.

In addition, we investigated the influence of the fragment order on the exam z-scores in the LA listening task. The exam z-scores represent the behavioral scores obtained based on the number of keywords that each person remembers correctly (for more information, see Eqlimi et al. (2020)). For this purpose, we used the likelihood ratio test (LRT) to compare the two following linear mixed-effect models:

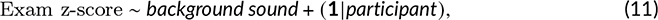

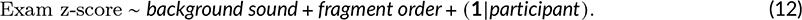

The corresponding *p*-value (Pr(*χ*^2^)) of LRT was *p* = 0.1038, suggesting that the predictor *fragment order* has no significant (linear) effect to model the exam z-scores.

### Link between exam results and T**_2_**

In our previous publication Eqlimi et al. (2020), a comprehensive statistical analysis was performed to relate behavioral results (exam scores) and frequency-domain characteristics of EEG signals. Here, with a similar methodology, we examine the relationship between exam results and T_2_ in the LA listening task. By comparing the model represented by formula 11 and the model in which T_2_ is added as a fixed predictor to model 4.11, we found no significant (linear) effect of T_2_ to improve the predictability of exam z-scores (*p*-value= 0.524). It is worth noting that this statistical analysis is based on time-averaged T_2_ and overall exam score over the presented fragment. The distribution of both exam scores and T_2_ over time is not necessarily linear, and the absence of a linear relationship between these variables might be explained to some extent.

### Time-varying complexity of microstates

Since recurrence time (T_2_) showed more statistical differentiation (refer to the previous subsection), the results of modeling the time series obtained from the epoch-wise (epochs of length 1 second) recurrence time using GAMM (formula 8) are shown in Figure 7. The significant results of smooth terms in the GAMM model to T_2_ show that (1) there was a significant random intercept (*F* = 1953.725, *edf* = 21.21, *p* < 2 × 10^−16^) and slope (*F* = 1058.335, *edf* = 18.90, *p* < 2 × 10^−16^) over *participant*.

**FIGURE 7.**
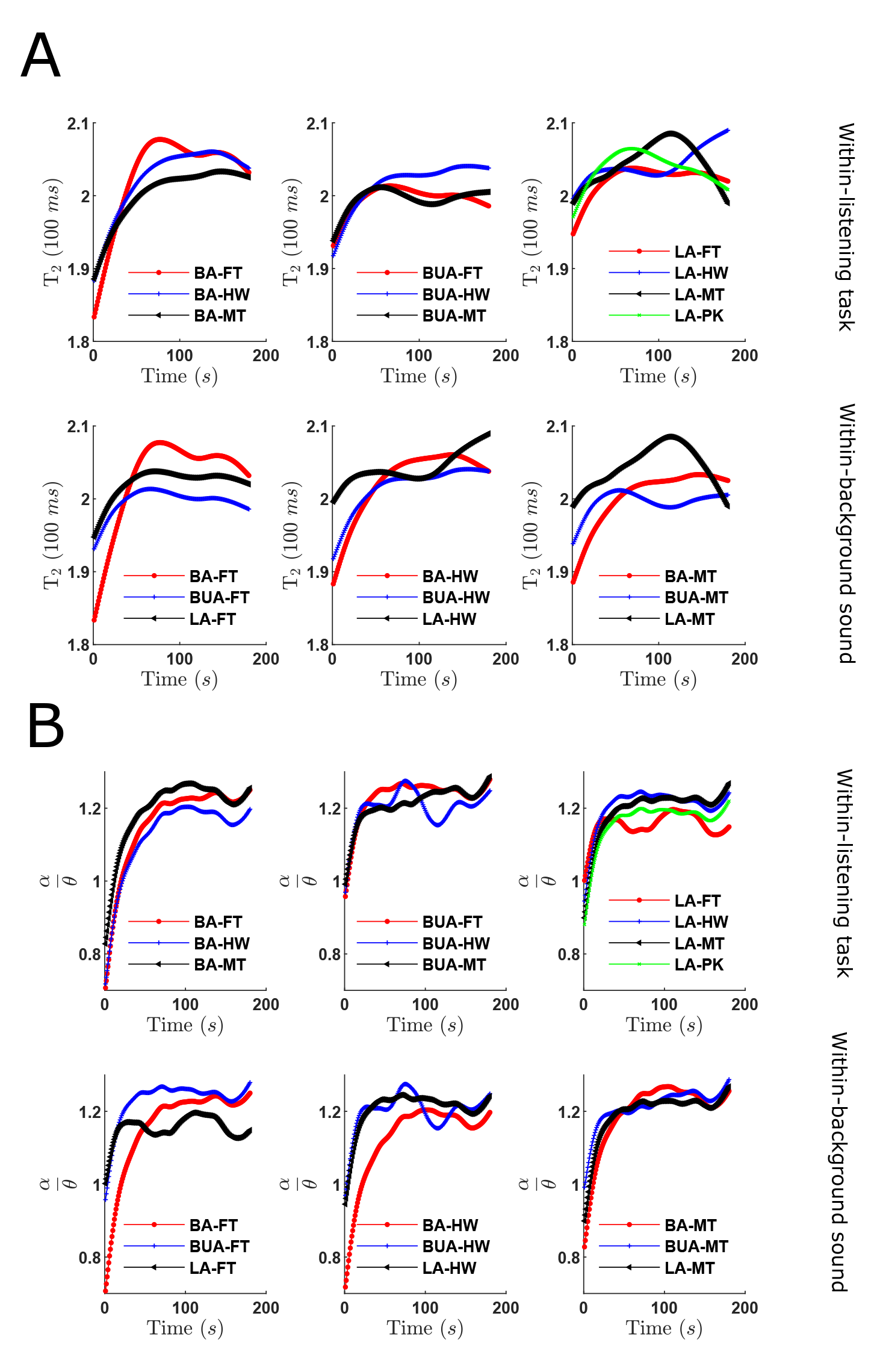
Time-varying T_2_ (panel **A**) and *α* /*θ* power ratio (averaged across all channels) (panel **B**) obtained using GAMM. The trends are colored by the listening conditions and represent the summed effect, i.e., the predicted response measure for a certain condition. Note that the random effect (*participant*) was excluded. The y-axis label in panel **A** represents a temporal resolution of 100 *ms* per increment. For example,T_2_ = 2 corresponds to a recurrence time of 200 *ms*.

Several studies (refer to section) have been done on EEG power spectral dynamics associated with listening. In our previous study (Eqlimi et al., 2020), we showed that PC_1_ (which is related to *α* power and *α* /*θ* power ratio) indicates a decrease in global attention. Motivated by this finding, we performed the time-frequency representation method to quantify the time-varying *α* /*θ* power ratio (averaged across all channels) and compare it with time-varying T_2_. The significant results of smooth terms in the GAMM model on the time-varying *α* /*θ* showed that there was a significant random intercept (*F* = 52563.364, *edf* = 21.946, *p* < 2 × 10^−16^) and significant random slope (*F* = 41991.249, *edf* = 20.078, *p* < 2 × 10^−16^) over *participant*.

Figure 7 illustrates the trends of time-varying T_2_ and *α* /*θ* power ratio, fitted using GAMMs, for all the listening conditions. Regardless of the listening condition, the figures visually indicate two notable patterns: (1) an initial rapid increase in T_2_ and *α* /*θ* within the first few tens of seconds and (2) a continuation of the trend with a lower slope thereafter, except for some conditions that exhibit a decreasing trend towards the end. Although T_2_ and *α* /*θ* show similar overall trends, there are distinct differences when comparing their responses across different situations. This suggests that T_2_ and *α* /*θ* may be independent. For example, in the multi-talker sound scenario, the T_2_ values for the LA task are higher compared to other tasks, while no such difference is observed for the *α* /*θ* ratio. Interestingly, despite the absence of significant mean differences in T_2_ between the background sounds, we found significant variations in T_2_ over time among the sounds. Specifically, when contrasting LA-PK (lecture attended in silence) and LA-FT, T_2_ exhibits significantly higher values (*p* < 0.05) within the time interval of [44 − 90] seconds in the LA-PK condition.

To compare time-varying T_2_ across the listening conditions, the difference curves in time between two specific listening conditions are shown in Figure 8 based on the GAMM predictions. To compare the conditions, separate GAMMS were fitted for each condition. The difference between their predicted values was calculated and smoothed for a visually clearer difference curve. Additionally, specific time intervals with significant differences between conditions were identified and marked with red lines on the smoothed difference curve. We conducted these comparisons for each background sound, enabling us to observe the differences between different listening tasks (BA vs BUA, LA vs BA, and LA vs BUA) in the same background noise. These results will be discussed in the subsection. It is worth mentioning that the random effect (inter-subject variability) is canceled out when the difference between the conditions is calculated. However, the size of the confidence intervals can be influenced by inter-subject variability.

**FIGURE 8.**
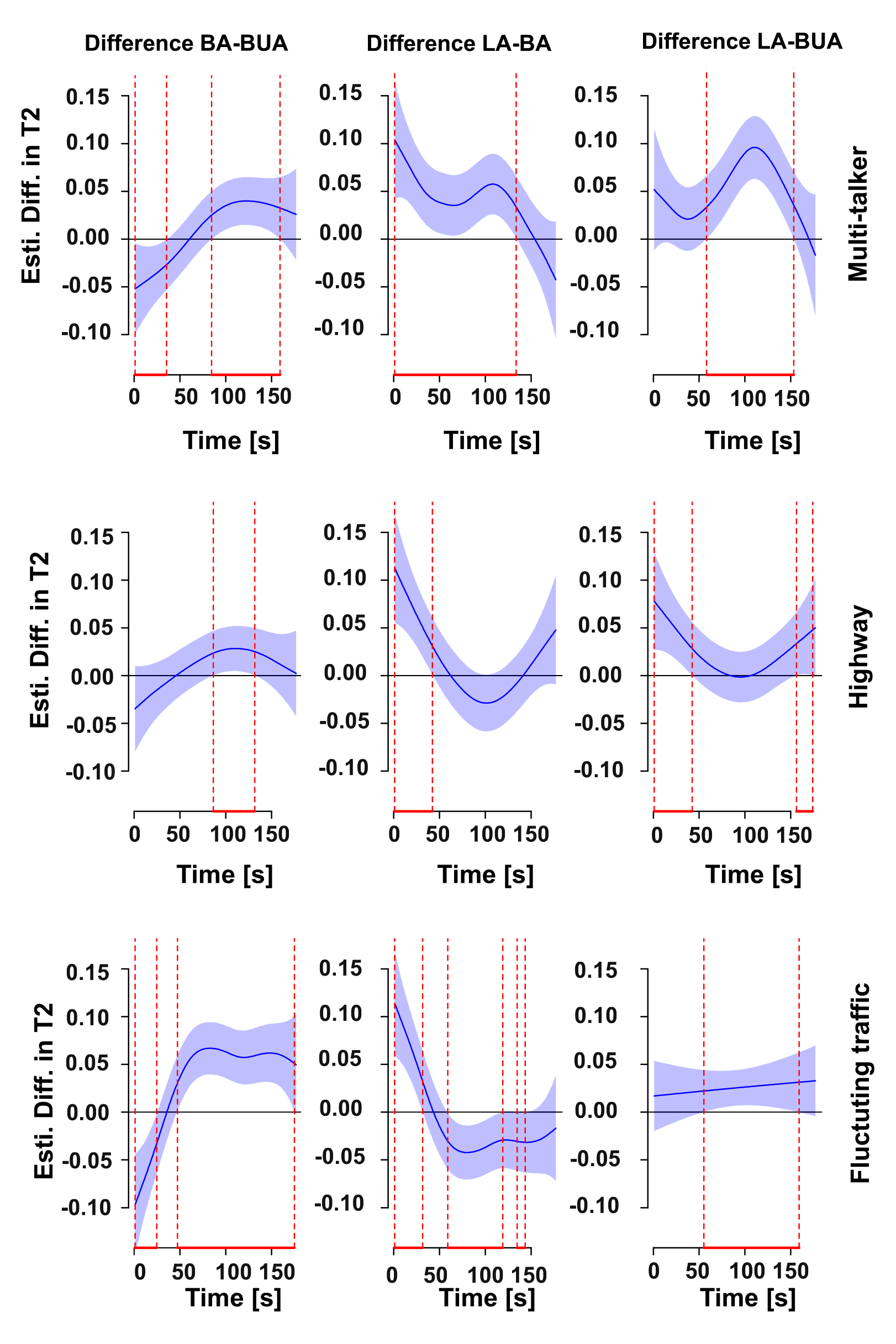
Estimated differences in microstate recurrence time (T_2_) between two listening tasks in a certain background sound. Significantly different time intervals between the given conditions are shown by red lines. The blue shaded areas indicate the 95% confidence intervals for the effect of time.

### Discussion and Conclusion

The present study aimed to investigate the temporal dynamics of time-resolved brain states and complexity analysis to understand the evolution of attentional engagement over time during speech perception in a noisy context. This paper is complementary to our previous research (Eqlimi et al., 2019, 2020), which focused on describing different listening conditions using spectral EEG components. This study reveals that the complexity of EEG microstate transitions is a pertinent measure for observing processes related to accumulating effort, attention switching, and memory during speech perception in noise. Although, the precise connection to memory recall remains unclear and requires the use of larger and more controlled datasets for a comprehensive understanding. Our primary measure of microstate complexity was the recurrence time, which refers to the duration it takes for a trajectory of a dynamical system to return to a previously visited neighborhood (Ngamga et al., 2012). A longer microstate recurrence time implies a greater duration for a microstate to recur in the brain.

Firstly, our findings showed that when participants focused their attention on speech in the presence of background noise (attended condition), the complexity of EEG microstate transitioning was significantly higher compared to the condition where they ignored the background noise sounds (unattended condition). Secondly, we showed the role of attention in modulating the duration of microstate C during speech perception in a noisy environment. Ignoring the background noise (unattended condition) led to a shorter duration of microstate C. Conversely, when attention was directed towards either speech or the background noise (attended conditions), the duration of microstate C was prolonged. Lastly, our findings revealed a distinct and consistent trend in the complexity of microstate transitioning over time (specifically, within a 3-minute duration) during all listening conditions, regardless of whether participants were attending to speech or the background noise. This trend can be divided into two distinct phases: an initial phase of lower complexity followed by a subsequent phase of increased complexity, eventually reaching a steady state. In sum, these findings highlight the crucial role of attention and memory encoding in shaping the complexity of brain states.

It is worth noting that our analysis in this study was based on single-trial EEG data, which distinguishes it from typical sensory stimulation paradigms that involve repeated specific stimuli. While the use of single-trial EEG data aligns well with ecological validity, it also poses certain challenges in terms of analysis.

### Microstate duration

Our results show that during LA, the duration of microstate A is significantly lower than during the other tasks, while the duration of microstate B is significantly higher. Milz et al. (Milz et al., 2016) obtained a similar trend while com-paring a verbalization to a visualization task. Actively listening to lectures could be regarded as a verbalization task or at least a task that evokes activity in the same brain regions, and hence our work confirms their findings. The seeming contradiction to earlier work (Britz et al., 2010) that suggests that in the resting state, microstate A is correlated with blood-oxygen-level-dependent (BOLD) imaging signal in the temporal cortex and associated with the auditory network, while microstate B is correlated with BOLD signal in occipital cortex and associated with the visual network, is explained in (Milz et al., 2016) by inhibition. EEG power in the alpha band is assumed to dominate and would reflect inhibition of the visual areas during verbalization and vice versa. During our LA task, alpha band power is indeed consistently higher than during the BA task, which could reflect inhibition, but this is not the case for the BUA task (see Table 1 in (Eqlimi et al., 2020)). It is worth noting that the interpretation by (Milz et al., 2016) is also criticized by (Custo et al., 2017). Microstate D duration is significantly higher during the BUA task than during the LA task. Microstate D is mostly related to the dorsal attention network (Britz et al., 2010; Seitzman et al., 2017). However, the authors in (Milz et al., 2016) propose that microstate D is more associated with focus switching and reorientation, hence, the occurrence of microstate D increases in the resting state compared to a goal-oriented task (Milz et al., 2017; Michel and Koenig, 2018). There is no significant difference between the duration of microstate D during the BA task compared to the BUA task, mainly because the distribution of duration widens during the BA task. This is probably because attentively listening to some background sounds (e.g. multi-talker babble) induces more focus switching than goal-oriented tasks used in other experiments and during LA. During all conditions in our experiment, the duration of microstate C is longer than that of any other state. It is however significantly lower during the BUA task. Microstate C (anterior-posterior orientation) has been linked to the cognitive control and salience network activation in the anterior cingulate-insula (Britz et al., 2010). The activity in the latter reflects maintained alertness, (Coste and Kleinschmidt, 2016) which is consistent with its increased duration during alertness-demanding tasks (LA and BA) compared to the resting state (BUA).

The duration of Microstate E is only significantly higher during the LA than during the BA. It has been suggested that microstate E in the resting state is likely related to the default mode network (DMN), where left mid frontal gyrus, anterior cingulate cortex (ACC), posterior cingulate cortex (PCC) are involved (Custo et al., 2017). The functional role of the left inferior frontal gyrus has been linked to the listening effort during the speech-in-noise task (Alain et al., 2018; Dimitrijevic et al., 2019; White and Langdon, 2021), which is in line with increased E duration during the LA compared to the BA. However, the lack of difference between the LA and BUA criticizes this interpretation. Since it has been suggested that different areas in the DMN act as hubs (Custo et al., 2017; Andrews-Hanna, 2012; Andrews-Hanna et al., 2010), restriction to a specific area will lead to misinterpretation.

Duration of microstate F (labeled as G in Custo et al. (2017)) is significantly lower during the BA task than during the other two tasks, i.e., the LA and BUA. Involvement of different areas in the appearance of this microstate has been shown, such as the right inferior parietal lobe and cerebellum Custo et al. (2017). Apart from the fact that the cerebellum plays a wide range of functional roles, especially in motor coordination, its role in language processing, regulating the acquisition of auditory sensory data, and working memory Desmond et al. (1997); Peterburs et al. (2021); Mariën and Borgatti (2018) has also been emphasized, which can be consistent with higher F duration during LA than during the BA. In addition, the inferior parietal lobe is also involved in the interpretation of sensory information and associating meanings to verbal information while listening to a narration (Saalasti et al., 2019).

### Microstates and Distraction

Previous research (Korn et al., 2021) revealed a decrease in microstate A and an increase in microstate C engagement when individuals focused on watching a video while being distracted by background music compared to the resting state. In our study, a pure resting state condition was not available. Instead, we used the BUA condition as a proximate substitute. Consistent with the findings of (Korn et al., 2021), our study showed that microstate C exhibited higher engagement during the LA compared to the BUA condition. This finding highlights the significance of microstate C in speech perception and the encoding of memory in the presence of distractors. Furthermore, we observed a lower engagement of microstate A during the LA condition compared to the BUA condition, aligning with the findings of (Korn et al., 2021). However, we also observed that microstates D and B played a significant role in distinguishing between the LA and BUA.

Recent research has revealed that high cognitive load, resulting from the interplay between task-relevant top-down processing and task-irrelevant bottom-up processing (distractors) in a perceptual decision-making task, affects microstate engagement (He et al., 2021). Specifically, it has been observed that there is an increase in the engagement of microstate C in the healthy control group, while microstates C and D exhibit heightened engagement in individuals with mild cognitive impairment (MCI). In the conditions of our study, the LA suggests a high cognitive load due to the involvement of top-down attention and memory processes, as well as the presence of distracting stimuli. The increase in microstates C and D observed in (He et al., 2021) aligns with our findings, where the LA condition exhibited the highest duration of microstate C compared to the BA and BUA conditions. In a related recent study (Kleinert et al., 2022), self-control, i.e., the ability to inhibit irrelevant stimuli and distractions was found to positively correlate with microstate duration, specifically microstate D, confirming our observation of increased microstate D during the LA.

### Microstate recurrence and complexity

In this work, we suggested a novel indicator based on the complexity quantification of the transitions that occur between different microstates. We hypothesized that these transitions reflect the listening state. By complexity, we mean the features that quantify the properties of microstate recurrence. Microstate recurrence features are estimated without taking into account Markovian and stationariness of neural transitions. We found that when subjects are listening to continuous speech (a lecture in silence or environmental noise in the LA), the recurrence time (T_2_) is higher than the other two listening tasks, i.e., the BA and BUA tasks. Moreover, the temporal features related to the recurrence of EEG microstates showed that, in the early stages of information processing, the dynamics and transitions between microstates were less complex (shorter T_2_), whereas more complex (longer T_2_) behavior was observed in the latter stages. In other words, the same region in microstate space is re-visited faster at first and gradually the recurrence speed decreases. Our findings are supported by the evidence presented in (Singer, 2013; Olguín-Rodríguez et al., 2018; Arzate-Mena et al., 2021), in which the non-stationary dynamical features and a kind of shadow of the evolution of an attractor in phase space are emphasized.

The transition in microstate behavior from lower complexity at the start of each fragment to higher complexity later on, or in other words, the increase in diversity of microstates visited during listening, may also be explained by a balance between sensory-driven and cortical processes. The cortical processes can be explained by the predictive coding model (Friston, 2009). In fact, the attractor in the microstate space, as the phase space of our neural dynamical system, might identify how much the brain uses its prior expectations to predict during listening. At the early moment of listening to a sound, we expect the brain to process the sensory signals. However, over time, the brain uses the predictions generated by our internal model of the world, which are fed back from top-down information. In other words, the accuracy of prior expectations may be reflected in the complexity of microstate transitioning. Strong evidence has been suggested in the literature on the relationship between the cortical microcircuits and functional networks and predictive coding (Bastos et al., 2012). During listening to a speech in noise, we might change our prediction to understand the sensory-driven information. In (Adams et al., 2021), the prediction error (especially in disorders like schizophrenia) is related to the uncertainty of our brain model about the world. They suggested increased confidence in sensory input and decreased confidence in the predictions, leading to a kind of uncertainty and delusions. Our results are consistent with this evidence, as the complexity of microstate transitions (which may reflect the uncertainty of our internal model) increases as the sensory input becomes more complex (speech vs background sounds). In addition, to interpret this evidence, they used an unstable attractor, which in our study is comparable to increasing recurrence time. Moreover, our results support the learning models of the orbitofrontal cortex (OFC) and norepinephrine neurons of the locus coeruleus (LC-NE) (Sadacca et al., 2017), where, OFC corresponds to a database of association and LC-NE plays a role in state creation.

### Alpha-to-Theta Power Ratio Time Course

Previous studies have placed considerable emphasis on spectral power, particularly theta and alpha power. In a study on episodic memory (Griffiths et al., 2021), it was found that alpha power decreases during the initial phase of information gathering, followed by an increase in theta power during the subsequent step of mnemonic binding. Based on our findings (refer to Figure 7, panel **B**), it becomes evident that during sensory processing in the earlier epochs of the alpha-to-theta power ratio time course, there is a noticeable decrease in alpha power, while theta power remains relatively constant (indicating a decrease in the alpha-to-theta power ratio). This observation suggests that as information is gathered and processed, there is a reduction in alpha activity. Our observed alpha suppression in the first phase aligns with the results and interpretation provided in the intracranial-EEG (iEEG) study (Nourski et al., 2021), which demonstrated the role of alpha suppression in regulating and modulating auditory processing in a speech perception task.

As mentioned in the later part of the time course, it is observed that there is a decrease in theta power (indicating an increase in the alpha-to-theta power ratio), which is associated with mnemonic binding. However, in our study, no binding was observed prior to the time interval in which a sentence is presented. Once binding starts to occur (which can be seen as running a predictive circuit and processing mismatch), there is a subsequent rise in alpha power. However, this increase is followed by a drop, indicating a complex interplay between information gathering and binding processes (refer to Figure 7, panel **B**). The decrease in theta power (resulting in an increase in the alpha-to-theta power ratio) observed in our findings is consistent with previous studies (Canales-Johnson et al., 2021; Griffiths et al., 2021; Köster et al., 2018; Pazienti et al., 2022). The previous studies support our findings by demonstrating that theta oscillations and slow waves are linked to an effortful top-down mnemonic reactivation in a mental imagery task (Canales-Johnson et al., 2021), in an episodic memory task (Griffiths et al., 2021), in an associative memory task (Koskinen et al., 2020), and in fading anesthesia (Pazienti et al., 2022).

In addition to our primary interpretation mentioned earlier, another possible explanation is the time-on-task effect. Previous studies have explored the relationship between alpha power evolution and this effect (e.g. (Benwell et al., 2019)). The increase in alpha power over time may imply a decrease in attention. However, the specific relationship between changes in alpha power and attention or vigilance has been debated (Wang et al., 2021b). Moreover, alpha activity is known to be associated with both endogenous and exogenous attention orientations, with these orientations being distinct from each other (Keefe and Störmer, 2021).

### Microstates and EEG Spectra link

There is an ongoing debate and no clear consensus regarding the link between EEG power spectra and microstates. A seminal study (Koenig et al., 2002) found a weak relationship between the time course of relative power values as a function of age and the time course of microstate characteristics as a function of age (correlation coefficient below 0.44). The relationship between spectra and microstates has been the subject of recent studies. For example, in (Croce et al., 2020), the role of the occipital alpha frequency band has been emphasized in mediating microstates in the resting state. However, it was also noted that the microstate switching is affected not only by alpha oscillations but also by interactions with other frequency bands. In a more recent study (Zulliger et al., 2022), it has been shown that a significant correlation exists between the time course of microstate features and the time course of power spectra in the delta, theta, alpha, and beta frequency bands, as observed in the within-subject analysis. Using between-subject analysis, significant correlations were found within the theta and alpha frequency bands. Qualitatively, our results align with the findings of (Zulliger et al., 2022), demonstrating a similar two-stage trend between the alpha-to-theta power ratio and the microstate complexity (*T*_2_) across time (starting with a low value in the early epoch and gradually increasing to a roughly stable level). Our interpretation is that the oscillatory activity, particularly in the theta and alpha frequency bands, plays a considerable role in shaping the transitions and complexity of microstates.

### EEG microstates and RSNs

At the beginning of the implementation of the EEG microstates, the attention of many researchers was to show the one-to-one correspondence between microstates and the RSNs (Musso et al., 2010; Khanna et al., 2015; Michel and Koenig, 2018), while recent research (e.g., (Antonova et al., 2022)) suggests that the correspondence is not that clear. As discussed in subsection, this complexity in the microstates-RSNs correspondence leads to identifying the functional significance of EEG microstates becoming challenging. From the very beginning of the emergence of the EEG microstates, the link between the microstates and RSNs was surprising because they are measured on different temporal scales (RSNs: 10 − 20 seconds and EEG microstates: 50 − 100 milliseconds (Britz et al., 2010; Creaser et al., 2021)). One of the well-established reasons for the microstates-RSNs link is that the EEG microstates time-series are scale-free, namely, they likely have a similar representation in different temporal scales (Van de Ville et al., 2010; Creaser et al., 2021). One of the ways to show the scale-free behavior of EEG microstates is long-range temporal correlations (LRTC) (the same method used for investigating the scale-free behavior of *α* oscillation in our previous publication (Eqlimi et al., 2020)). It has been suggested that EEG microstates have LRTC over six dyadic scales that span the two orders of magnitude (256 milliseconds to 16 seconds, which is comparable with the RSNs temporal scale) (Van de Ville et al., 2010). The presence of LRTC in EEG microstates, suggests that in addition to inherent characteristics of microstates, sequential connectivity of microstates may influence the transition between underlying sources (Britz et al., 2010; Creaser et al., 2021).

In this paper, two above-mentioned reasons, i.e., (1) the uncertainty in determining the functional role of each microstate and (2) the sequential connectivity, motivated us to focus more on transitions of the microstate sequences. For this purpose, the complexity quantification of EEG microstate transitions based on the recurrence quantification analysis is proposed and implemented, for the first time in this paper. One of the results of applying this method to the EEG data of this paper was that we were able to estimate the timescale of the recurrence of microstates, which is known as recurrence time. Our findings suggested that the recurrence time of microstates is an order of 80 − 210 milliseconds.

We showed that the 7 identified microstates, in this paper, are similar to microstates that are revealed in the resting-state literature. Although the similarity of topographic maps of resting-state and task-based EEG microstates has been shown in many studies (Seitzman et al., 2017; Gui et al., 2020; Zhang et al., 2022; Mheich et al., 2021), one of the important questions is how likely is it that these microstates are meaningful indicators of stimulus-related processing? This question is not accurately addressed in this paper, because we assume that the duration of microstates and transition complexity can be the indicators of task-related responses rather than the inherent spatial characteristics of microstates. Furthermore, in one of the recent studies (Antonova et al., 2022), the spatial similarity of the rest and task-based microstates was shown, and they used the duration of the microstates to differentiate different conditions, i.e. mind-wandering, verbalization, and visualization. They have shown the duration of microstate D is longer during mind-wandering than verbalization or visualization. Our findings suggest that the duration of microstate D during the BUA (background unattended) condition is longer than BA (background attended) and LA (lecture attended), which is in agreement with the results presented in (Antonova et al., 2022). Although the experimental tasks that have been performed in this study are not the same as the experimental tasks of this paper, mind wandering might frequently occur in the BUA condition in our experiment rather than the BA and LA conditions. In addition, we may expect more occurrences of verbalization and visualization in the LA compared to the BA and BUA. A detailed discussion has been presented in subsection. Altered resting-state EEG microstate dynamics have been shown in several neurodegenerative disorders, e.g. Alzheimer’s disease (Lian et al., 2021), schizophrenia da Cruz et al. (2020), and Parkinson’s disease Pal et al. (2021). The methodology performed in this paper may be used for identifying the cognitive dysfunctions in neurodegenerative disorders (for example, speech listening in Parkinson’s disease or schizophrenia).

### Microstate recurrence evolves differently over time depending on listening condition

Addressing one of our main questions, we asked whether there is any difference between the temporal dynamic of microstate complexity between the listening tasks (within-background-sound). Interestingly, our findings showed that this is indeed the case: despite the two-stage behavior described in the previous sections, temporal dynamics differ between the listening conditions. Comparing the time-varying T_2_ between the LA and BA (Figure 8, the middle panel) indicates that the microstate recurs slower in the LA at the early moment of listening, regardless of the type of the background sound (as we expected). This higher level of complexity reflects that the LA situation may require higher attention regulation and/or may solicit semantic processing and predictive coding more compared to the BA. This effect is also observed, in the middle and late moments of listening, for the background sounds of multi-talker and fluctuating traffic. Although we have linked the neural response of listening in the first few seconds to the processing of sensory information, attention switching occasionally happens between two external tasks or from an internal task to an external task (Verschooren, Sam, 2020). External attention refers to selecting the external task-relevant information and suppressing the task-irrelevant information, while internal attention is basically related to the information processing in the working memory (WM) (Oberauer, 2009; Chun et al., 2011). It has been shown that the attention switch can modulate ERP components (Verschooren et al., 2021). In our study, we expect that the LA task is more likely (compared to the BA) to cause attention switching (especially, between speech and background sounds). The high level of complexity in the early stages of information processing might be related to this mechanism.

The information content and the underlying semantic structures of lecture and multi-talker sound are similar. Therefore, during listening to the lecture in multi-talker sound, it might be more difficult to distinguish the target from the background sound which would be required switching attention back and forth between speech and background sound and between internal and external attention. This might be a reason for the longer time interval in Figure 8, where T_2_ is higher in the LA than the BA listening task in multi-talker sound.

The higher levels of complexity (in early moments) in the BUA compared to BA (Figure 8, the left panel, except the highway sound) can be also linked to the attention-switching effect. In the BA, the participants were supposed to focus on the background sounds, hence, the attention switch is less likely than in the BUA situation, where the participants were not supposed to pay attention to any sound, and their minds can wander. This might be a reason for the similarity between the LA and BUA tasks in the early moments (Figure 8, the right panel, except the highway sound). Although this similarity is due to a common effect between the LA and the BUA, that is, the participants do not focus continuously on the sensory input, the reason for this effect is different. In the LA, attention switching is likely due to semantic processing, while in the BUA, it is due to mind wandering and resting state. This difference is also seen in Figure 5, where the microstates that are visited are clearly different between the LA and the BUA in multi-talker sound.

In the later epochs of listening, internal attention emerges as the primary mechanism, leading to the involvement of working memory and predictive coding in information processing. Working memory is not solely limited to later epochs; it actually initiates from the first word we listen to. However, the crucial aspect is that as more information accumulates in working memory, the brain mechanisms responsible for its maintenance require additional cognitive resources to ensure successful processing. As a result, it is reasonable to expect higher levels of complexity in the comparisons between BA vs BUA and LA vs BUA and BA. However, there are instances where the presence of highway sounds contradicts this interpretation.

It is worth mentioning that T_2_ changes over time are strongly dependent on the background sound. However, for all the background sounds, initially, the BA is less complex (lower T_2_) as the people listen to the background sound or don’t listen to it, but they may still do some cognitive processing albeit not as much as for lectures where the message is dominant. The BUA also starts complex, but there is still an initial interest in the environment that grabs attention.

Although the main focus of this chapter was not to compare the response between different background sounds, it is worthwhile to mention that regarding the LA without the background sound (LA-PK), we found that (1) the initial high complexity, likely due to the linguistic processing, (2) the complexity increases over time, because more time might give more context for predictive coding, and (3) towards the end the complexity decreases due to mental fatigue and lack of motivation. In the LA-FT, initially, we see the same trend as LA-PK, but it exhibits lower complexity due to likely more time for the sensory processing needed. Towards the end, the complexity stays higher in the LA-FT as the predictive coding needs to intervene more, and the *α* /*θ* power ratio stays higher. Further studies are needed to statistically analyze as well as interpret the neural response differences between the background sounds.

### Data availability

Data available on reasonable request from the first author. The Matlab codes implementing the algorithms are publicly accessible on GitHub (https://github.com/EhsanEqlimi).

### Declaration of competing interest

The authors do not report any conflicts of interest.

### Credit authorship contribution statement

**Ehsan Eqlimi**: Conceptualization, Data curation, Formal analysis, Investigation, Methodology, Project administration, Software, Validation, Visualization, Writing-original draft, and Writing-review & editing. **Annelies Bockstael**: Conceptualization, Data curation, Funding acquisition, Supervision, Writing-review & editing **Marc Schönwiesner**: Concep-tualization, Funding acquisition, and Writing-review & editing. **Durk Talsma**: Conceptualization, Funding acquisition, and Writing-review & editing **Dick Botteldooren**: Conceptualization, Funding acquisition, Supervision, Project administration, and Writing-review & editing.

## Acknowledgment

This work was supported by “Flemish Government under the Onderzoeksprogramma AI Vlaanderen programme” and “Belgium Special Research Fund (BOF) of Ghent University”.

## Abbreviations

ACC: Anterior Cingulate Cortex
BA: Background Attended
BA-MT: Background Attended in Multitalker noise
BUA: Background Unattended
BUA-MT: Background Unattended in Multitalker noise
dBA: A-weighted Decibels
DMN: Default Mode Network
EEG: Electroencephalography
FT: Fluctuating Traffic noise
GAMM: Generalized Additive Mixed Model
GFP: Global Field Power
GMD: Global Map Dissimilarity
GLMM: Generalized Linear Mixed Model
HW: Highway noise
iEEG: Intracranial Electroencephalography
KL: Krzanowski-Lai criterion
LA: Lecture Attended
LA-FT: Lecture Attended in Fluctuating Traffic noise
LA-HW: Lecture Attended in Highway noise
LA-MT: Lecture Attended in multitalker noise
LA-PK: Lecture Attended in Pink noise
LC-NE: Locus Coeruleus-Norepinephrine
LOI: Line of Interest
MCI: Mild Cognitive Impairment
MT: Multitalker Noise
MRI: Magnetic Resonance Imaging
OAE: Otoacoustic Emission
OFC: Orbitofrontal Cortex
PCC: Posterior Cingulate Cortex
PK: Pink noise
RQA: Recurrence Quantification Analysis
RSN: Resting State Network
TFR: Time-Frequency Representation
t-SNE: t-Distributed Stochastic Neighbor Embedding
WM: Working Memory

