## Supplementary text for "Time course of EEG complexity reflects attentional engagement during listening to speech in noise"

#### 1 Modified k-means algorithm for EEG microstate analysis

The method used to identify EEG microstates in this paper is based on the modified k-means clustering algorithm introduced in [1]. This appendix describes the steps of this algorithm.

Assuming $n_{ch}$ EEG channels and $n_{ms}$ underlying microstates, the microstate model can be represented in vector form as follows:

$$\mathbf{x}(t) = \sum_{q=1}^{n_{ms}} \mathbf{a}_q s_q(t) + \mathbf{e}(t) \quad t = 1, \ldots, n_t, \tag{1}$$

where $\mathbf{x}(t) \in \mathbb{R}^{n_{ch} \times 1}$ includes the scalp electric potential measurements at time instant $t$, $\mathbf{a}_q \in \mathbb{R}^{n_{ms} \times 1}$ represents the $q^{th}$ microstate (normalized scalp topography), $s_q(t) \in \mathbb{R}^{1 \times 1}$ is the $q^{th}$ microstate intensity at time instant $t$, and $\mathbf{e}(t) \in \mathbb{R}^{n_{ch} \times 1}$ represents the zero-mean additive noise (independent and identically distributed for all time instants).

Assuming that only one microstate is active at any time instant, all $s_q(t)$ must be zero except for one (i.e., matrix $\mathbf{S}$ is 1-sparse). In the literature, this assumption is called the non-overlapping microstates or the discreteness assumption. The modified k-means algorithm tries to estimate the microstates ($\mathbf{a}_q$) based on this assumption. However, in [2], it is shown that microstates and their transitions are more continuous than discrete fashion, i.e., more than one microstate is active at any given moment. Part of this problem might be addressed by defining the microstates using the GFP peaks, but it is still not possible to guarantee that only one microstate is active even at the peak points. Some algorithms, called k-sparse component analysis ($k$-SCA), have tried to solve a similar problem when more than one source (here, microstate) is active at each time instant [3, 4, 5].

The non-overlapping microstates assumption (source disjointedness in $k$-SCA) causes the following constraint, which makes solving the problem (i.e., estimating microstates) easier:

$$\begin{aligned} s_{q_1}(t)\ .\ s_{q_2}(t) = 0\ ,\forall\ q_1 \neq q_2, \forall\ t \\ \sum_{q=1}^{N_{ms}} s_q(t)^2 \geq 1, \forall\ t \end{aligned} \tag{2}$$

The modified k-means clustering method aims to minimize the following cost functions with respect to all $a_q$ and $s_{kt}$, under constraints 2.

$$\mathcal{F} = \sum_{t=1}^{t=n_t} ||x(t) - \sum_{q=1}^{q=n_{ms}} a_q s_q(t)||_2, \tag{3}$$

where $\|\bullet\|_2$ stands for $\ell_2$-norm.

The modified k-means clustering method finds local minima of $\mathcal{F}$ using an iterative approach. It starts by randomly choosing $n_{ms}$ topographic maps from EEG data. In the first iteration, the spatial dissimilarity between these $n_{ms}$ topographic maps and every topographic map (EEG measurement vectors) is calculated. Various criteria can be used for this purpose, such as orthogonal squared distance (equation 4) and global map dissimilarity between each measurement vector and each microstate.

$$d_q(t)^2 = \mathbf{x}(t)^{\mathrm{T}}.\mathbf{x}(t) - (\mathbf{x}(t)^{\mathrm{T}}.\mathbf{a}_q)^2 \tag{4}$$

Each measurement ($\mathbf{x}(t)$) is labeled as belonging to the closest microstate, i.e.:

$$\hat{\mathcal{L}}(t) = \arg\min_x\ d_q(t)^2, \tag{5}$$

where $\hat{\mathcal{L}}(t)$ denotes the microstate label in time instant $t$.

The microstate intensity ($\hat{s}_k(t)$) can be estimated by:

$$\hat{s}_k(t) = \mathbf{x}(t)^{\mathrm{T}}\ .\ \mathbf{a}_k, \tag{6}$$

where $k = \hat{\mathcal{L}}(t)$.

The steps 4-6 are performed for all $\mathbf{x}(t)\ \forall\ t$. The goal of the next step is to estimate the microstates. i.e., $\mathbf{a}_q$. For this purpose, given the microstates labels ($\mathcal{L}(t)$), the minimum of $\mathcal{F}$ can be estimated by the normalized eigenvector corresponding to the largest eigenvalue of the matrix:

$$\mathbf{R}_q = \sum_{t/\mathcal{L}(t)=q} \mathbf{x}(t)^{\mathrm{T}}.\mathbf{x}(t), \tag{7}$$

$t/\mathcal{L}(t) = q$ represents only time points for which $\mathcal{L}(t) = q$.

Therefore, we can solve the following problem using the eigenvector decomposition (EVD):

$$\hat{\mathbf{a}}_q = \arg\max_{\mathbf{v}} \mathbf{v}^{\mathrm{T}}\mathbf{R}\mathbf{v} \ \textit{subject to} \ ||\mathbf{v}||_2 = 1 \tag{8}$$

The pseudo-code and the details on the algorithm convergence have been reported in [1].

### 2 Recurrence quantification analysis

This supplementary text aims to explain how the complexity features are calculated based on recurrence quantification analysis (RQA) in the context of transition complexity analysis of EEG microstates. The concepts of phase space, recurrence, and recurrence plot (RP) have been explained in the main manuscript. Here the RQA features, i.e., recurrence rate (RR), determinism (DET), trapping time (TT), laminarity (LAM), entropy (ENT), and recurrence time ($\mathrm{T}_2$) are formulated.

A thresholded recurrence plot (RP) can be obtained based on the pairwise Euclidean distance between the microstate global map dissimilarity (GMD) vectors (as phase vectors) as:

$$\mathrm{R}_{ij}^{n_{ms},\epsilon} = \Theta(\epsilon - ||y_i - y_j||_2), \quad y_i, y_j \in \mathbb{R}^{n_{ms}\times 1}, \quad i, j = 1, ..., n_t, \tag{9}$$

where $n_{ms}$ is the number of microstates (the dimension of phase space), $n_t$ is the number of time points (with the step of 100 milliseconds), $\Theta(\bullet)$ is the Heaviside function, $\epsilon$ is a threshold distance, $y_i$, and $y_j$ are the GMD values at time $i$ and $j$, respectively. The RQA features for each sub-matrix of RP, $\hat{\mathbf{R}}$, can be calculated as follows:

$$\mathrm{RR}^{\epsilon,n} = \frac{1}{n^2 - n}\sum_{i\neq j=1}^{n} \hat{\mathrm{r}}_{i,j}, \tag{10}$$

$$\mathrm{DET}^{\epsilon,n} = \frac{\sum_{l=d_{min}}^{n} lH_d^{\epsilon}(l)}{\sum_{i,j=1}^{n} \hat{\mathrm{r}}_{i,j}}, \tag{11}$$

$$\mathrm{TT}^{\epsilon,n} = \frac{\sum_{l=v_{min}}^{n} vH_v^{\epsilon}(l)}{\sum_{l=v_{min}}^{n} H_v^{\epsilon}}, \tag{12}$$

$$\mathrm{LAM}^{\epsilon,n} = \frac{\sum_{l=v_{min}}^{n} vH_v^{\epsilon}(l)}{\sum_{i,j=1}^{n} \hat{\mathrm{r}}_{i,j}}, \tag{13}$$

$$\mathrm{ENT}^{\epsilon,n} = -\sum_{l=d_{min}}^{n} \frac{H_d(l)}{\sum_{l=d_{min}}^{n} H_d(l)} ln(l), \tag{14}$$

$$\mathrm{T}_2^{\epsilon,n} = \frac{\sum_{w=1}^{n} wH_w^{\epsilon}(w)}{\sum_{w=1}^{n} H_w^{\epsilon}}, \tag{15}$$

where $\hat{r}_{i,j}$ is $\{i, j\}$ element of $\hat{\mathbf{R}}$, $n$ is the number of time points in $\hat{\mathbf{R}}$, $l$ is the length of black lines in $\hat{\mathrm{R}}$, $H_d$, $H_V$ are the histogram of the lengths of the diagonal and vertical structures, respectively, $d_{min}$, and $v_{min}$ are the minimum thresholds for considering the diagonal and vertical lines, respectively, $w$ and $H_w$ are the length and histogram of white vertical lines, respectively.

The descriptive specifications of these RQA features are presented in Table 1.

Table 1: Descriptive specifications of RQA features. Note that the black color for $\mathbf{r}_{ij} = 1$ or recurrence points and white color for $\mathbf{r}_{ij} = 0$ have been used.

| RQA feature | RP characteristics | Description |
|---|---|---|
| RR | Black dots | Relative density of recurrence points |
| DET | Black diagonal lines | Repeating recurrences (diagonal structures) |
| TT | Black vertical lines | How long the state is trapped |
| LAM | Black vertical lines | Repeating recurrences (vertical structures) |
| ENT | Black diagonal lines | Complexity of the deterministic structure |
| $T_2$ | White vertical lines | Time needed to recur |

### 3 EEG data quality control and preprocessing

Since the EEG data acquisition process was done continuously during consecutive listening conditions, the probability of the presence of artifacts and noise can increase over time due to possible tiredness and more movement. However, after the end of the second condition (background attended, BA) and before the written exam, participants stretched their arms and legs. Although they still participated in the exam with the presence of EEG electrodes on their heads, the electrode connections were checked again.

As described in the main manuscript, the preprocessing procedure was performed separately for each condition (LA, BA, and BUA). Then, for each condition, the continuous EEG data was divided into fragments based on the trigger channels corresponding to the presented sounds. Here, we aim to quantify the quality of EEG data across the sequential conditions, in the sense of how different artifacts change over time. For this purpose, the IClabel algorithm [6] was applied to the components obtained from the continuous EEG data (per condition) using infomax independent component analysis (ICA).

IClabel tries to classify the components in the classes *brain*, *muscle*, *eye*, *heart*, *channel noise* and *others* based on their spatial topography. As explained in our previous publication [7], only the components related to the eye have been removed. Here, we aim to apply the ICLabel algorithm to the results of the first ICA (before removing eye-related components).

For each class, the average of the IClabel classification probabilities over the fragments presented in each listening condition is calculated. Figure 1 displays the distribution of these classification probabilities over the whole test population. The higher overlap between the distributions of each subplot means that the probability of the class corresponding to this subplot is less affected by the passage of time (equivalently, by the listening conditions; order of presentation:1-LA, 2-BA, and 3-BUA).

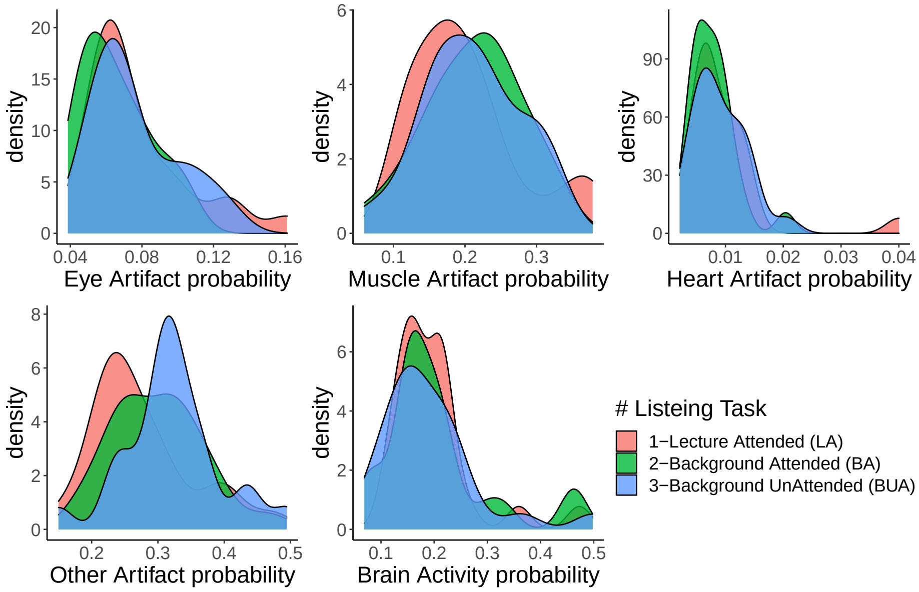


Figure 1: Density plot: the probability of IClabel classes grouped by the sequential listening task. The probability of different types of artifacts (eye, muscle, heart, and others) and brain activity probability, in the 64-channel EEG data obtained in this thesis, were estimated based on IClabel [6] algorithm, and then it was averaged over the presented fragments in listening conditions. The density plots represent the distribution of these average probabilities over the whole test population. The higher overlap between the distributions of each subplot means that the probability of the class corresponding to this subplot is less affected by the passage of time (equivalently, by the listening conditions; order of presentation:1-LA, 2-BA, and 3-BUA).

### 4 Inter-subject EEG microstate variability

In the main manuscript, it was explained how to identify the microstate topographic maps from the aggregated EEG data of each subject. We aggregated all the EEG signals obtained in different fragments of different listening conditions, and then microstate analysis was applied. The result of this analysis was the seven microstate topographic maps for each subject. Then, to compute the grand average microstate maps across subjects, the extracted topographic maps were permuted within each subject in a way that maximized the commonality across cases. In this appendix, we intend to investigate the inter-subject variability of these microstate topographic maps. For this purpose, a 3-dim multidimensional scaling (MDS) plot of identified 64-dim microstate topographic maps for all subjects is shown in Figure 2. In addition, the resulting individual microstate maps are illustrated in Figures 3, 4, and 5.

These results show that although the spatial similarity between the grand average topographies and topographies of most of the subjects is acceptable, there are also some individual topographies with noticeable differences (likely due to local and unwanted activities). In this thesis, the distance (GMD) between the topographies of each subject and the grand average microstate topographies (along the fragments presented) is considered as the main feature. As a result, if there are activities (whether caused by unwanted factors or task-related activities), the given microstate will be less pronounced. However, the inter-subject microstate variability requires considerably more investigation in both resting-state and task-based EEG data.

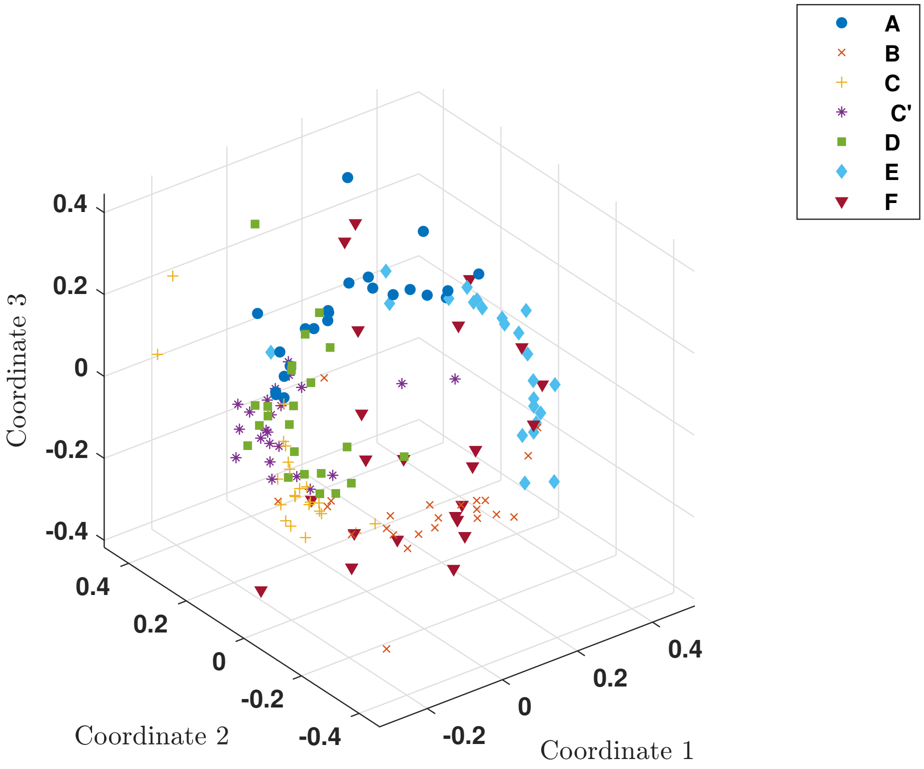


Figure 2: 3-dim multidimensional scaling (MDS) plot of identified 64-dim microstate topographic maps for all subjects. Colored by microstate classes (A-F and C′). Each point corresponds to one of the seven microstate topographic maps for one of the 23 subjects.

### References


[1] Roberto D Pascual-Marqui, Christoph M Michel, and Dietrich Lehmann. Segmentation of brain electrical activity into microstates: model estimation

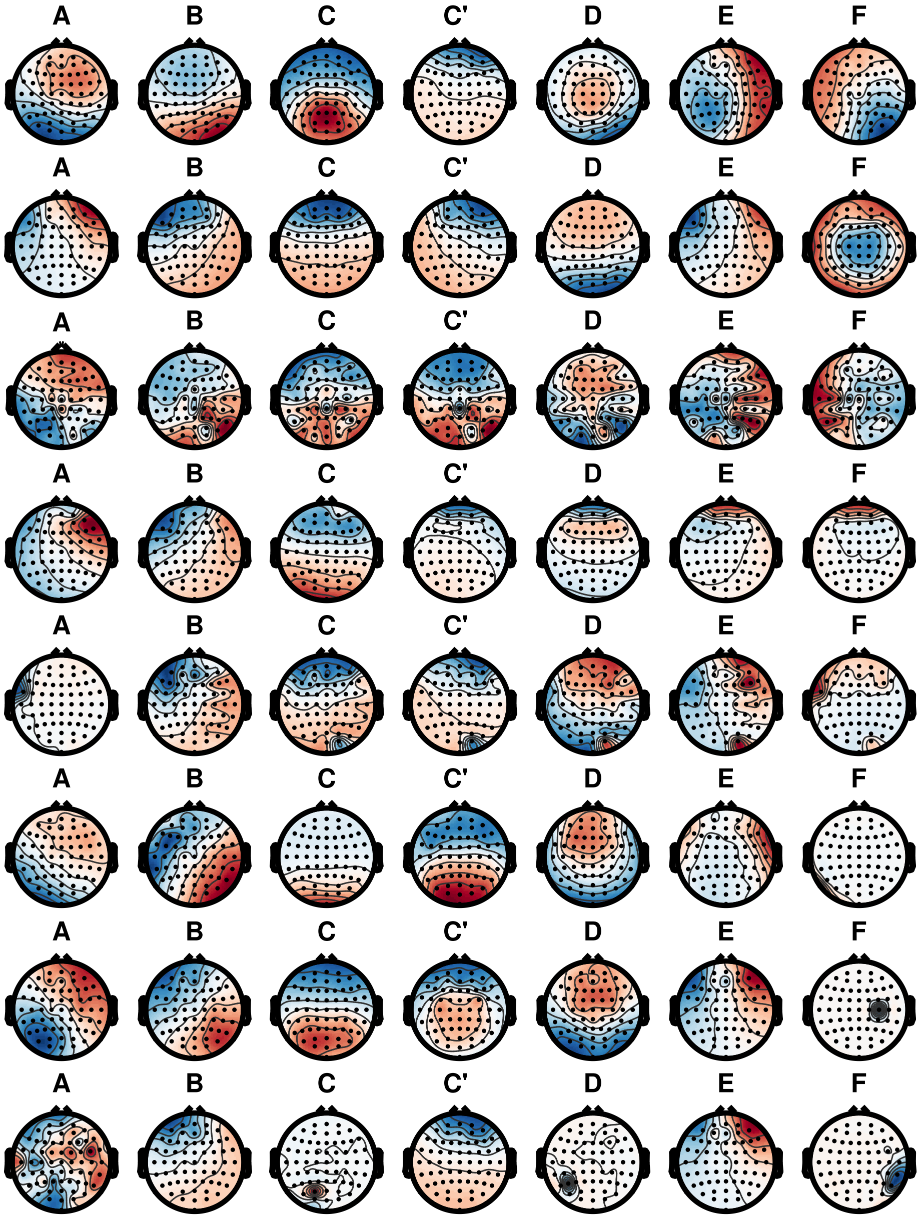


Figure 3: Microstate topographic maps identified from subject No. 1-8.

and validation. *IEEE Transactions on Biomedical Engineering*, 42(7):658–665, 1995.

[2] Ashutosh Mishra, Bernhard Englitz, and Michael X Cohen. EEG microstates

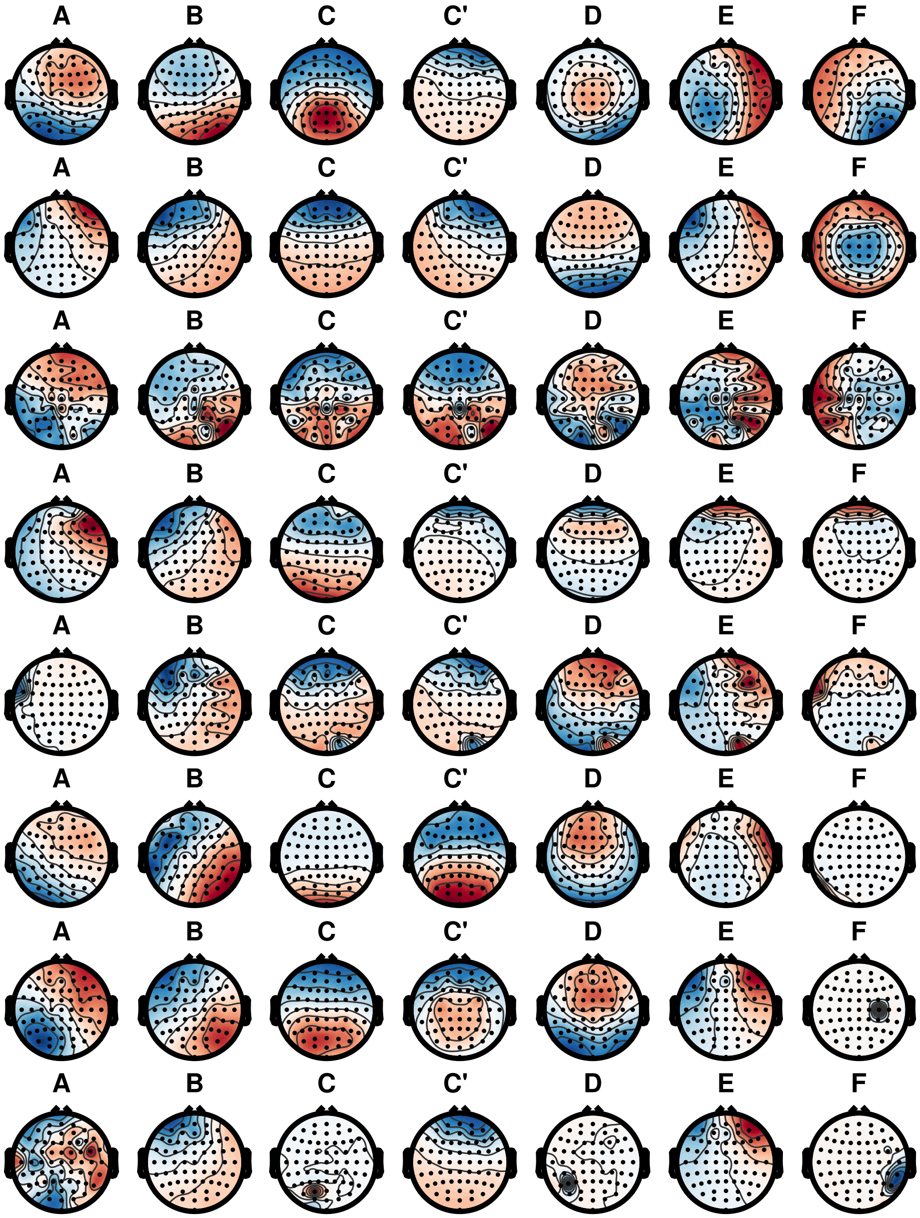


Figure 4: Microstate topographic maps identified from subject No. 9-16.

as a continuous phenomenon. *NeuroImage*, 208:116454, 2020.

[3] Ehsan Eqlimi and Bahador Makkiabadi. Multiple sparse component analysis based on subspace selective search algorithm. In *2015 23rd Iranian*

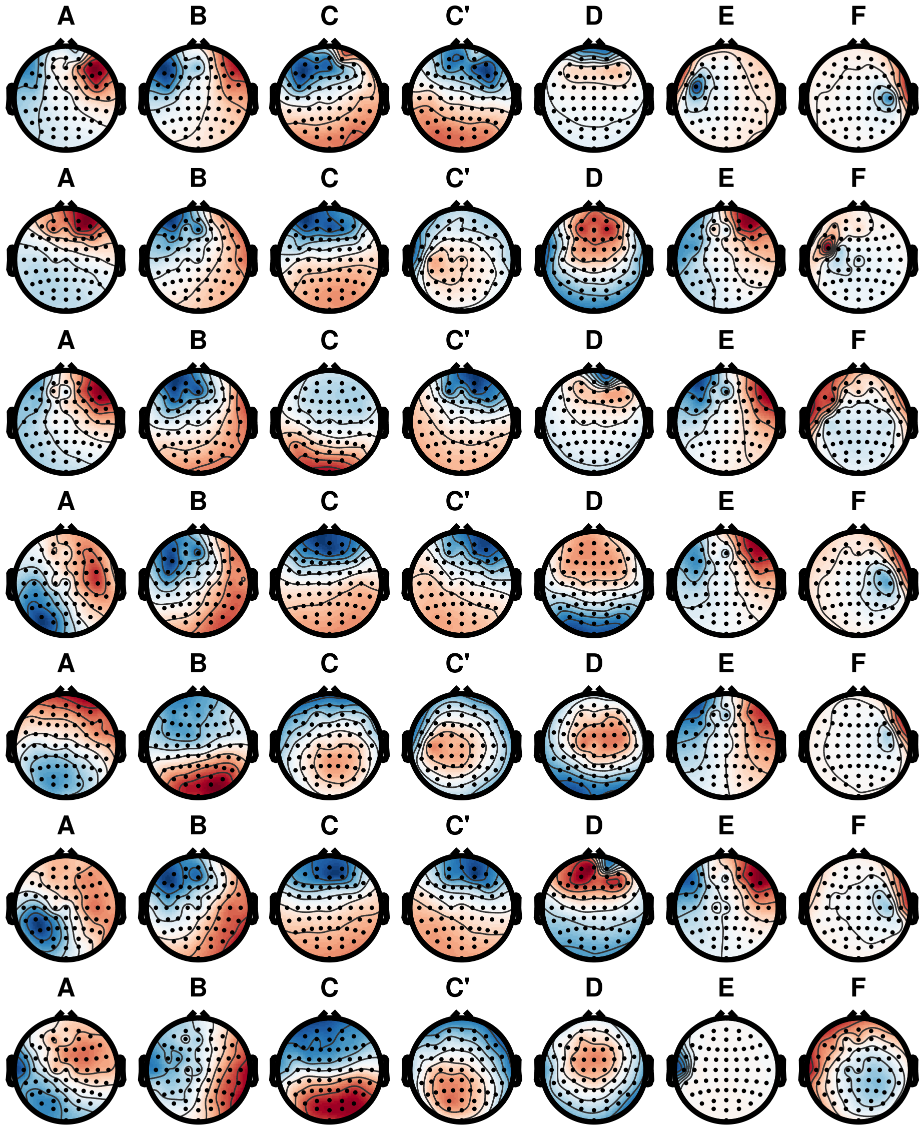


Figure 5: Microstate topographic maps identified from subject No. 17-23.

*Conference on Electrical Engineering*, pages 550–554. IEEE, 2015.

[4] Ehsan Eqlimi, Bahador Makkiabadi, Nasser Samadzadehaghdam, Hassan Khajehpour, Fahimeh Mohagheghian, and Saeid Sanei. A novel underdetermined source recovery algorithm based on k-sparse component analysis. *Circuits, Systems, and Signal Processing*, 38(3):1264–1286, 2019.

[5] Ehsan Eqlimi, Bahador Makkiabadi, Ardeshir Fotouhi, and Saeid Sanei.

Underdetermined blind identification for $k$-sparse component analysis using ransac-based orthogonal subspace search. *arXiv preprint arXiv:2008.03739*, 2020.

[6] Luca Pion-Tonachini, Ken Kreutz-Delgado, and Scott Makeig. Iclabel: An automated electroencephalographic independent component classifier, dataset, and website. *NeuroImage*, 198:181–197, 2019.

[7] Ehsan Eqlimi, Annelies Bockstael, Bert De Coensel, Marc Schönwiesner, Durk Talsma, and Dick Botteldooren. EEG correlates of learning from speech presented in environmental noise. *Frontiers in Psychology*, 11, 2020.
